# TEA-seq: a trimodal assay for integrated single cell measurement of transcription, epitopes, and chromatin accessibility

**DOI:** 10.1101/2020.09.04.283887

**Authors:** Elliott Swanson, Cara Lord, Julian Reading, Alexander T. Heubeck, Adam K. Savage, Richard Green, Xiao-jun Li, Troy R. Torgerson, Thomas F. Bumol, Lucas T. Graybuck, Peter J. Skene

**Affiliations:** Allen Institute for Immunology, Seattle, WA, USA; GlaxoSmithKline, Collegeville, PA, USA; Department of Pediatrics, University of Washington, Seattle, WA, USA

## Abstract

Single-cell measurements of cellular characteristics have been instrumental in understanding the heterogeneous pathways that drive differentiation, cellular responses to extracellular signals, and human disease states. scATAC-seq has been particularly challenging due to the large size of the human genome and processing artefacts resulting from DNA damage that are an inherent source of background signal. Downstream analysis and integration of scATAC-seq with other single-cell assays is complicated by the lack of clear phenotypic information linking chromatin state and cell type. Using the heterogeneous mixture of cells in human peripheral blood as a test case, we developed a novel scATAC-seq workflow that increases the signal-to-noise ratio and allows simultaneous measurement of cell surface markers: Integrated Cellular Indexing of Chromatin Landscape and Epitopes (ICICLE-seq). We extended this approach using a droplet-based multiomics platform to develop a trimodal assay to simultaneously measure <u>T</u>ranscriptomic state (scRNA-seq), cell surface <u>E</u>pitopes, and chromatin <u>A</u>ccessibility (scATAC-seq) from thousands of single cells, which we term TEA-seq. Together, these multimodal single-cell assays provide a novel toolkit to identify type-specific gene regulation and expression grounded in phenotypically defined cell types.

## Introduction

Peripheral blood mononuclear cells (PBMCs) purified using gradient centrifugation are a major source of clinically relevant cells for the study of human immune health and disease (Böyum, 1968). Like most other human tissues, PBMCs are a complex, heterogeneous mixture of cell types derived from common stem cell progenitors (Laurenti and Göttgens, 2018). Despite the genome being mostly invariant between different PBMC cell types, each immune cell type performs an important and distinct function. Understanding the genomic regulatory landscape that controls lineage specification, cellular maturation, activation state, and functional diversity in response to intra- and extracellular signals is key to understanding the immune system in both health and disease (Satpathy et al., 2019; Wang et al., 2020; Zheng et al., 2020).

Recent improvements in single-cell genomic methods have enabled profiling of the regulatory chromatin landscape of complex cell type mixtures. In particular, droplet-based single-cell assays for transposase-accessible chromatin (scATAC-seq, dscATAC-seq, mtscATAC-seq) allow profiling of open chromatin at single-cell resolution (Buenrostro et al., 2015; Lareau et al., 2019). Promising new methods have combined scATAC-seq with simultaneous measurement of nuclear mRNAs (e.g. sci-CAR, Cao et al., 2018; SNARE-seq, Chen et al., 2019; SHARE-seq, Ma et al., 2020). However, identification of highly specified functional immune cell types is hampered by current computational labeling and label transfer methods, in part due to complexity in linking regulatory sites to gene expression. To overcome these limitations, we systematically tested whole cell and nuclear purification and preparation methods for PBMCs. We found that intact, permeabilized cells perform extremely well for scATAC-seq, exceeding conventional scATAC-seq on nuclei by some measures (**Figure 1b**). This insight enables a new protocol analogous to Cellular Indexing of Transcriptomes and Epitopes (CITE-seq; Stoeckius et al., 2017) to measure both surface protein abundance and chromatin accessibility: Integrated Cellular Indexing of Chromatin Landscape and Epitopes (ICICLE-seq, **Figure 1a** and **Figure 3**). Finally, we demonstrate that our optimized permeable cell approach can be combined with a droplet-based multiomics platform to enables the simultaneous measurement of three different molecular compartments of the cell: mRNA (by scRNA-seq), protein (using oligo-tagged antibodies), and DNA (by scATAC-seq), which we term TEA-seq after <u>T</u>ranscription, <u>E</u>pitopes, and <u>A</u>ccessibility (**Figure 4**). This new assay provides a complete picture of the molecular underpinnings of gene regulation and expression at the single cell level.

**Figure 1.**
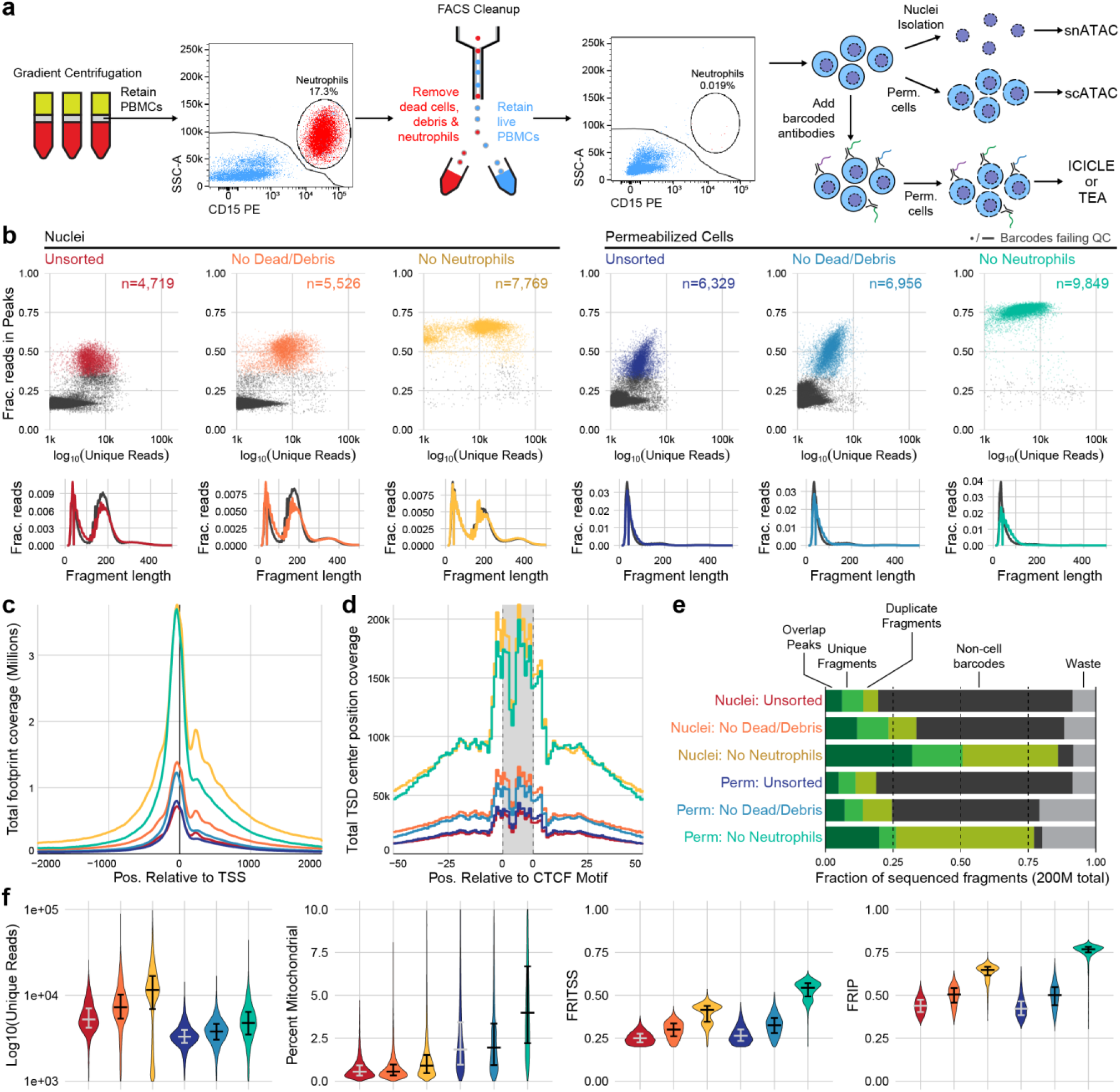
**a**, Schematic overview of major steps in snATAC, scATAC, and ICICLE-seq methods. **b**, Comparison of quality control characteristics of ATAC-seq libraries generated from nuclei isolation and permeabilized cells, with and without FACS. Top panels show signal-to-noise as assessed by fraction of reads in peaks (FRIP) on the y-axis, and quantity of unique fragments per cell barcode on the x-axis. Lower panels display fragment length distributions obtained from paired-end sequencing of ATAC libraries. Colored lines represent barcodes that pass QC filters; gray lines represent barcodes failing QC (non-cell barcodes). All libraries were equally downsampled to 200M total sequenced reads for comparison. Colors in b are re-used in remaining panels. **c**, Total coverage of Tn5 footprints summed across all transcription start sites (TSS). Tn5 footprints are 29 bp regions comprising the 9 bp target-site duplication (TSD) and 10 bp on either side, which represent accessible chromatin for each transposition event. **d**, Total coverage of TSD centers summed over a set of 100,000 genomic CTCF motifs found in previously published DNase hypersensitive sites(Meuleman et al., 2020). TSD centers are obtained by shifting +4 and −5 bp from the 5’ and 3’ ends of uniquely aligned fragments, respectively. **e**, Barplot representations of the fraction of total aligned reads in various QC categories. Fragments overlapping a previously published peak set for PBMC dscATAC-seq(Lareau et al., 2019) are in the ‘Overlap Peaks’ category. Unique fragments are the remaining uniquely aligned fragments that do not overlap peak regions. ‘Waste’ reads were not aligned or were assigned to cell barcodes with fewer than 1,000 total reads. **f**, Violin plots showing distributions of QC metrics. Median (wide bar) and 25th and 75th quantiles (whiskers and narrow bars) are overlaid on violin plots. Median values are also in **Table 1**. Note that the y-axis of the first panel is on a logarithmic scale; remaining panels are linear.

**Table 1.** QC metrics summary for experiments displayed in **Fig. 1**. Median % Mitochondrial was calculated as a fraction of total fragments; FRITSS and FRIP were calculated as a fraction of unique fragments. Neut., neutrophils; FRITSS, fraction of reads in transcription start sites; FRIP, fraction of reads in peaks.

| Source Type | FACS Depletion | N Nuclei/Cells<br>Pass QC | Median<br>N Fragments | Median<br>% Mitochondrial | Median<br>N Unique | Median<br>FRITSS | Median<br>FRIP |
| --- | --- | --- | --- | --- | --- | --- | --- |
| Nuclei | Unsorted | 4719 | 7344 | 0.6% | 5247 | 0.249 | 0.439 |
| Nuclei | Dead/Debris | 5526 | 10526 | 0.6% | 7284 | 0.301 | 0.505 |
| Nuclei | Dead/Debris/Neut. | 7769 | 19972 | 0.9% | 11528 | 0.415 | 0.648 |
| Perm. Cells | Unsorted | 6329 | 5541 | 1.9% | 3308 | 0.264 | 0.422 |
| Perm. Cells | Dead/Debris | 6956 | 6733.5 | 2.0% | 3795.5 | 0.326 | 0.502 |
| Perm. Cells | Dead/Debris/Neut. | 9849 | 14069 | 4.0% | 4756 | 0.544 | 0.768 |

## Results

### Optimization of single-nucleus and single-cell ATAC-seq of PBMCs

Our initial scATAC-seq experiments followed the protocol described by 10x Genomics, which largely adhered to the Omni-ATAC workflow (Corces et al., 2017). This protocol utilizes a combination of hypotonic lysis, detergents, and a saponin to isolate nuclei without releasing mitochondrial DNA. After performing this assay, sequencing, and tabulating data quality metrics (**Methods**), we identified two major populations of cell barcodes (**Figure 1b**, left panel): (1) A large number of barcodes, shown in gray, that have a low number of unique fragments and a low Fraction of Reads in Peaks (FRIP). These barcodes contain little useful information but consume 80% of total sequenced reads (**Figure 1e**, non-cell barcodes) at a sequencing depth of 200 million reads per library (20,000 reads per expected barcode); (2) Barcodes with higher quality as measured by FRIP (red points) that contain enough information to attempt downstream analysis.

The loss of 80% of sequenced reads to non-cell barcodes is costly. Previous studies of scRNA-seq data have shown that cellular lysis can release ambient RNA that increases the abundance of low-quality barcodes and contaminates droplets, yielding barcodes with both cellular and ambient RNAs that reduces the accuracy of the transcriptional readout (Marquina-Sanchez et al., 2020). We reasoned that nuclear isolation protocols may cause the release of ambient DNA, causing a similar effect in scATAC-seq datasets. Optimization of nuclear lysis protocols, especially changing to less stringent detergents, provided increased FRIP and decreased non-cell barcodes (**Figure 1 - figure supplement 1** and **Figure 1 - figure supplement 2a**). Hypotonic lysis conditions used in these protocols may also be a biophysical stressor to the native chromatin state, as previously observed (Lima et al., 2018). To reduce perturbation of chromatin, we performed cell membrane permeabilization under isotonic conditions to allow access to the nuclear DNA without isolating nuclei through hypotonic lysis. The saponin digitonin was used to cause concentration-dependent selective permeabilization of cholesterol-containing membranes while leaving inner mitochondrial membranes intact, preventing high levels of Tn5 transposition in mitochondrial DNA (Adam et al., 1990; Colbeau et al., 1971). Digitonin has previously been used for ATAC-seq assays under hypotonic conditions in Fast-ATAC (Corces et al., 2016) and plate-based scATAC-seq (Xi Chen et al., 2018) protocols. Permeabilization of intact cells under isotonic conditions greatly reduced the amount of non-cell barcodes and their contribution to sequencing libraries (**Figure 1 - figure supplement 2b**).

### Removal of neutrophils greatly increases scATAC-seq data quality

We also observed that PBMCs purified by leukapheresis rather than Ficoll gradient centrifugation had consistently higher FRIP scores and fewer non-cell barcodes (**Figure 1 - figure supplement 2c**). A major difference between our Ficoll-purified PBMCs and these leukapheresis-purified PBMCs was the presence of residual neutrophils in our Ficoll-purified samples. We tested removal of dead cells and debris with and without removal of neutrophils using Fluorescent Activated Cell Sorting (FACS) from PBMC samples with high neutrophil content (**Figure 1a** and **Figure 1 - figure supplement 3**). When applied to either nuclei (**Figure 1b**, left panels) or permeabilized cells (**Figure 1b**, right panels), there was a large increase in FRIP and reduction in non-cell barcodes in our scATAC-seq libraries (**Figure 1e**). Removal of neutrophils did not have an adverse effect on leukapheresis-purified PBMCs (**Figure 1 - figure supplement 1c**, right panel), and depletion using anti-CD15 magnetic beads also improved data quality (**Figure 1 - figure supplement 2d** and **Figure 1 - figure supplement 4**), though not to the same extent as FACS-based depletion. Staining and flow cytometry using an 8-antibody panel (**Supplementary Table 1**) on Ficoll- and leukapheresis-purified PBMCs showed minimal effect of the magnetic bead treatment on non-neutrophil cell type abundance (**Supplementary Table 2**).

### Comparing single-cell and single-nucleus ATAC characteristics

We assessed the quantitative and qualitative differences between nuclei and permeabilized cell protocols with and without sorting by performing both protocols on a single set of input cells. Permeabilized cells yielded many more high-quality cell barcodes than nuclear preps using equal loading of cells or nuclei (15,000 loaded, expected 10,000 captured, **Table 1**). scATAC-seq libraries obtained from nuclei had many more reads with fragments originating from nucleosomal DNA fragments (**Figure 1b**, lower panels), and non-cell barcodes from nuclei (gray lines) contained more of these fragments than cell barcodes. Thus, an overabundance of mononucleosomal fragments may indicate non-cell fragment contamination. Libraries from permeabilized cells consisted almost entirely of short fragments, suggesting that permeabilization under isotonic conditions did not loosen or release native chromatin structure at the time of tagmentation (**Figure 1b**, lower panels). Previous bulk ATAC-seq studies have shown that differing nuclear isolation protocols lead to varying amounts of mononucleosomal fragments (Li et al., 2019). In agreement with *in vitro* experiments studying the effects of low salt on nucleosomal arrays (Allahverdi et al., 2015), this further suggests that hypotonic lysis leads to alteration of chromatin structure, raising the possibility of artifactual measurements of accessibility in nuclei-based ATAC-seq. To assess the effect that this difference has on the data obtained by each method, we overlaid Tn5 footprints near transcription start sites (TSS, **Figure 1c**) and CTCF transcription factor binding sites (TFBS, **Figure 1d**). The signal at TSS was retained in permeabilized cells, but positions flanking the TSS (occupied by neighboring nucleosomes) had reduced signal compared to isolated nuclei (examined in detail in **Figure 1 - figure supplement 5**). At CTCF motifs, we observed nearly identical patterns of accessibility in both nuclei and permeabilized cells, suggesting that scATAC-seq signal at regulatory TFBS is retained in permeabilized cells. Overall, permeabilized intact cells obtained by FACS had the highest FRIP and fraction of reads in TSS (FRITSS) scores, fewest non-cell barcodes, and greatest cell capture efficiency with only a modest increase in mitochondrial reads (**Table 1, Figure 1e,f**).

### Improved label transfer and differential analysis

We next examined the effect of methodological differences on downstream biological analyses (**Figure 2**). Removal of neutrophils greatly improved the ability to separate various cell types in UMAP projections of both nuclei and cells (**Figure 2a-b**). To provide ground truth for label transfer, we performed flow cytometry on an aliquot of the same cell sample used for scATAC-seq, above. A panel of 25 antibodies (**Supplementary Table 3**) was used to determine the proportion of each of the 12 cell types used to label the scATAC-seq cells in the PBMC sample (**Figure 2 - figure supplement 1** and **Supplementary Table 4**). Label transfer was enabled by the ArchR package (Granja et al., 2020) to generate gene scores and perform transfer from a reference scRNA-seq dataset using the label transfer method provided in the Seurat package (Stuart et al., 2019) (**Methods**). Using these tools, removal of neutrophils improved label transfer scores, and permeabilized cells yielded more cells with high label transfer scores than nuclei-based approaches (**Figure 2b-c**). In addition, permeabilized cells provided labels most similar to the cell type proportions identified by flow cytometry (**Figure 2d**), with identification of CD8 effector cells only observed in scATAC-seq with permeabilized cells. All methods yielded fewer CD16+ monocytes than observed by flow cytometry, suggesting that CD16+ monocytes may be lost during scATAC-seq using either nuclei or permeabilized cells, or that label transfer methods were not conducive to identifying this cell type (**Figure 2d**). After labeling cell types, we used ArchR to call peaks for each cell type and perform pairwise tests of differential accessibility between each pair of cell types (**Figure 2e**). We found many more differentially accessible sites in both cells and nuclei after removal of neutrophils. Differential accessibility was also used to identify differentially enriched TFBS motifs in each cell type (**Figure 2f**). Without neutrophil removal (Nuclei Unsorted, top panel), we were unable to identify significantly enriched motifs in B cells and NK cells that were readily apparent in data from clean nuclei or permeabilized cells (bottom two panels). Together, these results demonstrate that neutrophil removal and the use of permeabilized cells allow for identification of specific cell types and TFBS motifs that are involved in regulation of gene expression.

**Figure 2.**
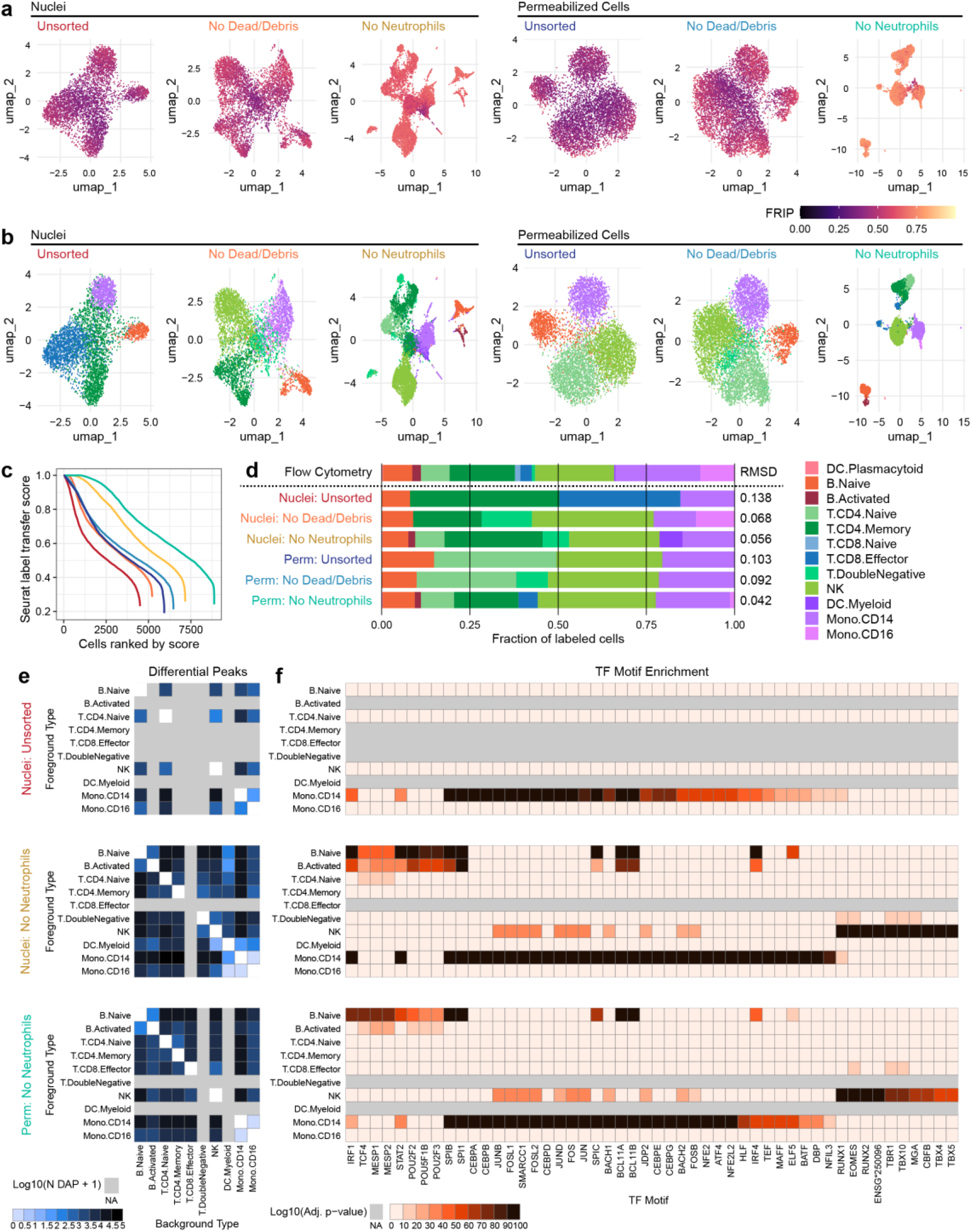
**a**, UMAP projection plots for corresponding datasets in Figure 1. Points are colored based on a common scale of fraction of reads in peaks (FRIP), bottom right. The number of cells in each panel are displayed in Figure 1b. **b**, UMAP projection plots colored based on cell type obtained by label transfer from scRNA-seq (**Methods**). Colors for cell types are below, to the right. **c**, To visualize the number and quality of transferred labels, we ranked all cells based on the Seurat label transfer score obtained from label transfer results, and plotted lines through the score (y-axis) vs rank (x-axis) values. **d**, Bar plot showing the fraction of cells in each dataset that were assigned each cell type label. The top row shows cell type proportions for the same PBMC sample obtained by 25-color immunotyping flow cytometry (**Methods, Supplementary Table 3, and Figure 2 – figure supplement 1**). Root-mean-square deviation (RMSD) values were computed by comparison of labelled cell type proportions to values derived from flow cytometry (**Supplementary Table 4**). Colors for cell types are to the right of the barplot. **e**, Heatmap plots of pairwise differentially accessible peaks (DAPs) between each pair of types, as computed with Wilcoxon tests using the ArchR package(Granja et al., 2020) (**Methods**). Values shown are the number of peaks more highly accessible in the foreground type (y-axis) compared to the background type (x-axis). Colors represent Log10-scaled and binned counts of DAPs, as shown in the scale to the bottom-left. Gray regions represent cell types that were not observed in a dataset. **f**, Heatmap plots of enriched transcription factor binding site (TFBS) motifs. Colors represent −1 x Log10-scaled and binned adjusted p-values for tests of enrichment (**Methods**). Values > 100 are colored black.

### Joint measurement of chromatin accessibility and surface epitopes with ICICLE-seq

Under standard scATAC-seq protocols, removal of the cell membrane severs the connection between the cell surface and the chromatin state of cells. By retaining the cell surface on permeabilized cells, we were able to extend scATAC-seq to simultaneously profile cell surface proteins and chromatin accessibility, which we term Integrated Cellular Indexing of Chromatin Landscapes and Epitopes (ICICLE-seq, **Figure 3** and **Methods**). The ICICLE-seq protocol utilizes a custom Tn5 transposome complex with capture sequences compatible with the 10x Genomics 3’ scRNA-seq gel bead capture reaction for simultaneous capture of ATAC fragments and polyadenylated antibody barcode sequences (**Figure 3 - figure supplement 1** and **Supplementary Table 5**). Antibody-derived tags (ADTs) from oligo-antibody conjugates (**Supplementary Table 6**) and ATAC-seq fragments can then be selectively amplified by PCR to generate separate libraries for sequencing (**Figure 3 - figure supplement 1**). Due to the nature of fragment capture in this system, we obtain both a cell barcode and a single-end scATAC-seq read. We performed ICICLE-seq on a leukapheresis-purified PBMC sample using a 46-antibody panel, and were able to obtain 10,227 single cells with both scATAC-seq and ADT data from three capture wells that passed adjusted QC criteria: > 500 unique ATAC fragments (median = 761), FRIP > 0.65 (median = 0.725). Cells passing ATAC QC had a median of 3,871 ADT UMIs per cell (**Figure 3 - figure supplement 2b**). UMAP projection and ATAC label transfer on ICICLE-seq data had resolution similar to scATAC-seq on intact permeabilized cells after dead cell and debris removal (**Figure 3b**). We were able to leverage the additional ADT data to cluster and identify cell types based on their cell surface antigens (**Figure 3c-e**) at higher resolution and accuracy (**Figure 3f**). UMAP based on ADT data and Jaccard-Louvain clustering allowed identification of cell type-specific clusters (**Figure 3d**) based on clear association of cell type-specific markers with clusters (**Figure 3e** and **Figure 3 - figure supplement 2**). Once identified, we leveraged these cell type labels to identify differentially accessible peaks (DAPs) in the scATAC-seq data (**Figure 3g**), even for types that were not separated based on label transfer from scRNA-seq (e.g. exhausted T cell subtypes). Thus, ICICLE-seq provides a novel platform for the identification of cell types in scATAC-seq data based on well-established cell surface markers.

**Figure 3.**
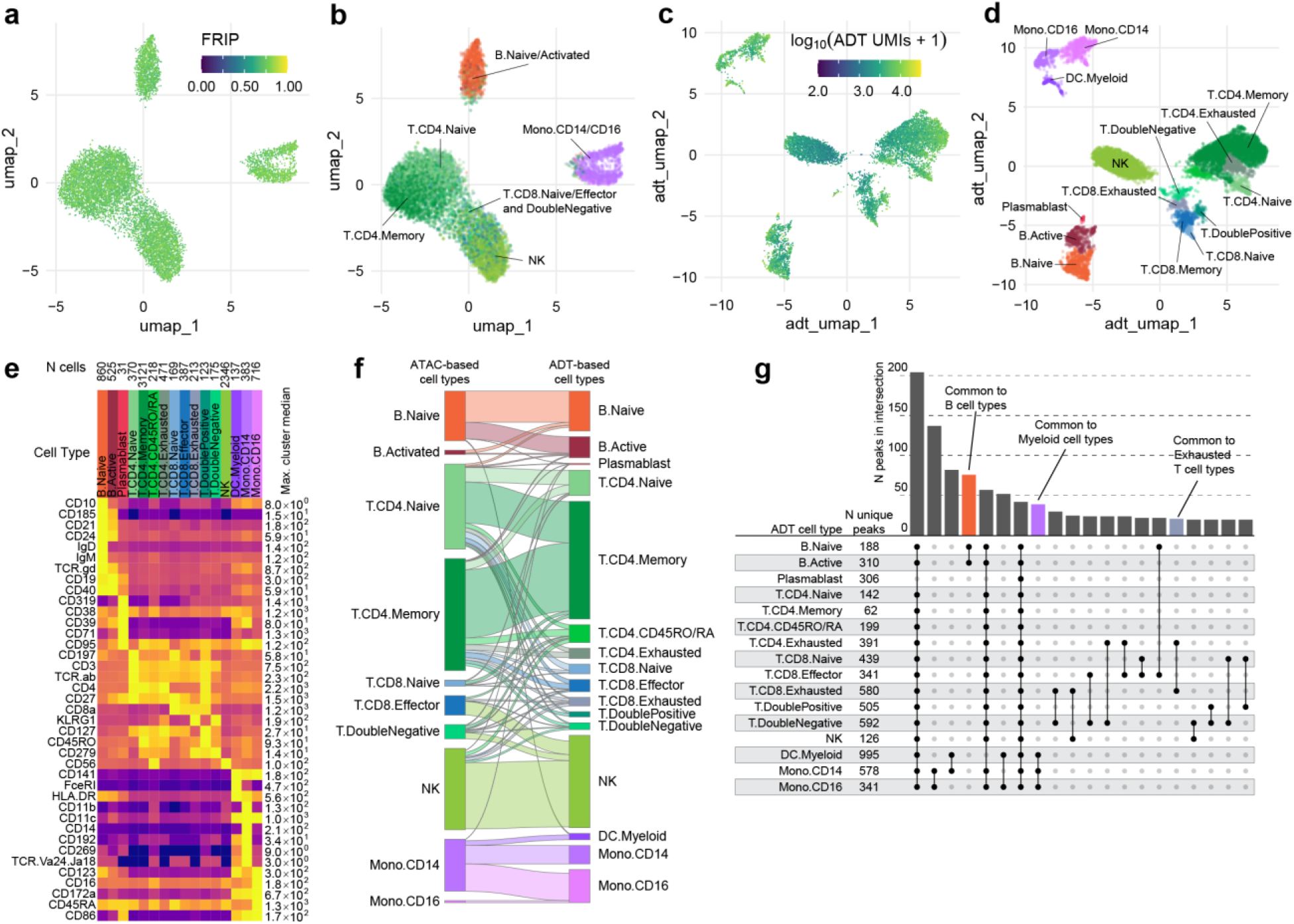
**a**, UMAP projection plot of ICICLE-seq cells based on scATAC-seq data. Cells are colored based on fraction of reads in peaks (FRIP). **b**, UMAP projection of scATAC-seq data, as in **a**. Cells are colored based on cell type labels obtained by ArchR label transfer (**Methods**). **c**, UMAP projection plot of ICICLE-seq cells based on antibody-derived tag (ADT) data. Cells are colored according to the total number of unique molecule indexes (UMIs) across all markers. **d**, UMAP projection based on ADT data, as in **c**, colored according to cell type labels derived from marker expression (**Methods**). **e**, Heatmap of median ADT count values for each marker in each cell type labeled in panel d. Values are separately scaled in each row between zero and the maximum value (right column) for each marker. **f**, Alluvial plot showing the relationship between ATAC-based cell type labels derived from label transfer in **b** and cell type labels derived from ADT-based markers in panel **d**. **g**, Peaks were called on aggregated single-cell data for each ADT-based cell type. For each cell type, the top 2,500 peaks were compared (except for plasmablasts, for which 592 peaks were identified). The number of unique peaks is displayed to the right of each cell type label, and peaks found in 20 combinations of cell types are displayed to the right. Closed/black points indicate cell types that are members of each intersection set, and bars above these points show the number of peaks in each intersection.

### Trimodal measurement of transcripts, epitopes, and accessibility with TEA-seq

With the release of a commercially-available platform for simultaneous capture of RNA-seq and ATAC-seq from single nuclei, we reasoned that unfixed, permeabilized cells could be used to perform simultaneous capture of three major molecular compartments: DNA can be captured using scATAC-seq, RNA could be captured using scRNA-seq, and protein epitope abundance can be captured using polyadenylated antibody barcodes, which we term TEA-seq after <u>T</u>ranscripts, <u>E</u>pitopes, and <u>A</u>ccessibility. After trials and optimization of key steps, we were able to obtain libraries on the 10x Genomics Multiome ATAC plus Gene Expression platform that combined all three of these measurements for thousands of single cells (**Figure 4a,b**) using a panel of 46 oligo-tagged antibodies (**Supp Table 6**). After initial data processing, we identified 7,939 cell barcodes that passed the QC criteria for scATAC-seq described above (median = 8,655 Unique ATAC fragments, **Figure 4c**) and had > 750 genes detected by scRNA-seq (median = 3,543 RNA UMIs; median = 1,617 genes detected, **Figure 4d**). Additional QC metrics are provided in **Figure 4 – figure supplement 1**.

**Figure 4.**
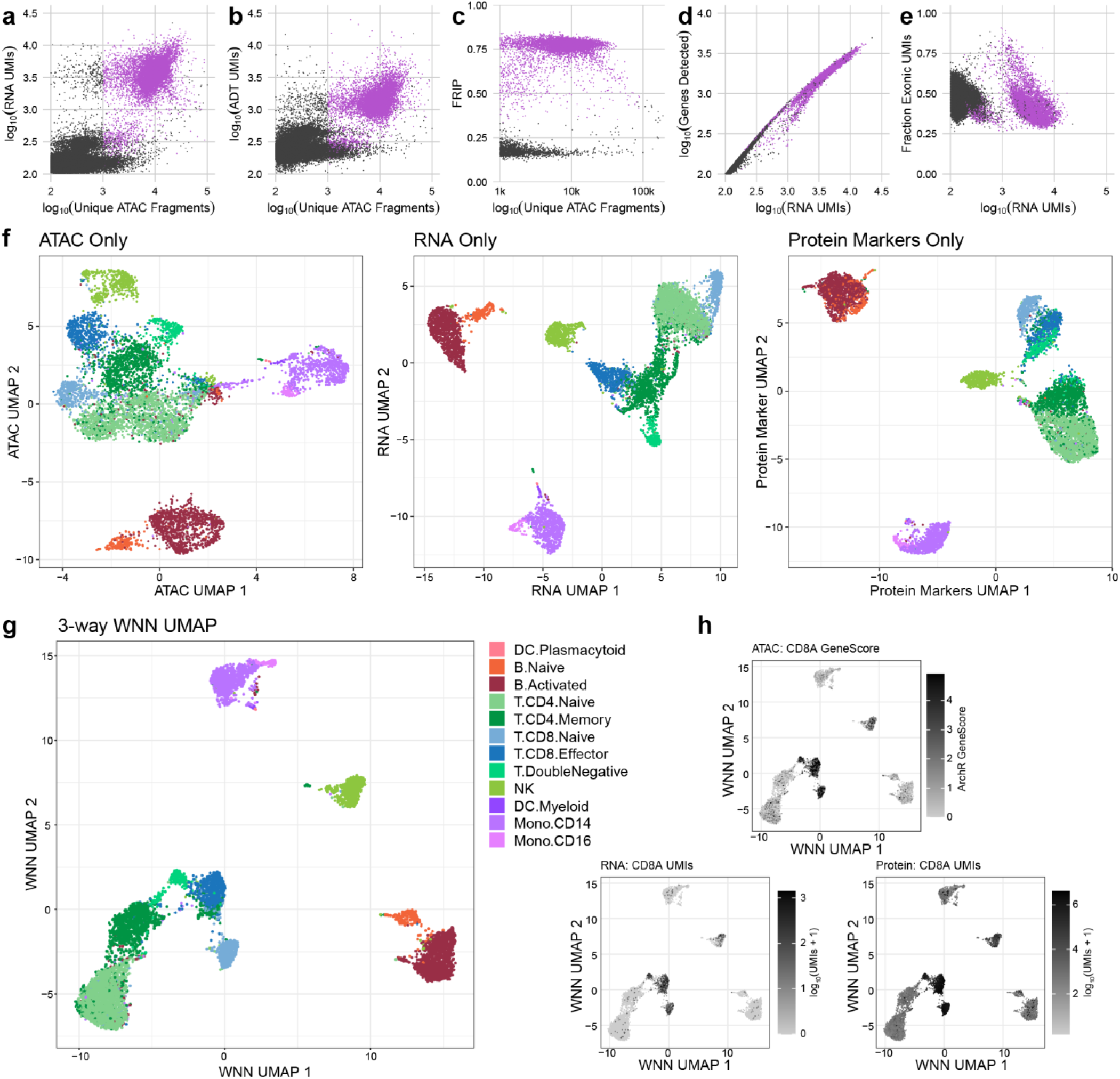
**a**, Scatterplot comparing unique scATAC-seq fragments and scRNA-seq UMIs for each TEA-seq cell barcode. In panels a-e, n = 61,546 barcodes are displayed in total; 7,939 passing QC criteria are represented by purple points. **b,** Scatterplot comparing unique scATAC-seq fragments and ADT UMIs for each cell barcode. **c,** scATAC-seq QC scatterplot comparing unique scATAC-seq fragments and FRIP scores for each cell barcode. **d**, scRNA-seq QC scatterplot comparing the number of scRNA-seq UMIs to the number of genes detected for each cell barcode. **e,** Comparison of the number of scRNA-seq UMIs to the fraction of scRNA-seq reads derived from exons. **f,** UMAP projections generated using each of the three modalities separately. **g,** A joint UMAP projection generated using 3-way weighted nearest neighbors (WNN) that leverages all three of the measured modalities. **h,** Detection of a single target, CD8A, across all 3 modalities. For scATAC-seq, GeneScore values generated using the ArchR package are displayed; for scRNA-seq and Protein/ADT, the number of UMIs detected for the CD8A gene and anti-CD8a antibody are used, respectively.

The alignment method used for these datasets allowed us to quantify the number of RNA UMIs that aligned to intronic vs exonic regions of the transcriptome. We found that reads mapped to both introns and exons (median 39.2% exonic, **Figure 4e** and **Figure 4 – figure supplement 1e**), suggesting capture of a combination of cytoplasmic mRNAs and intron-containing nuclear transcripts. These cell barcodes also had a median of 1,311 ADT UMIs per cell, enabling us to leverage cell surface epitopes to infer functional states of each cell. For each modality, we were able to perform dimensionality reduction and UMAP projection to get separate views of the relationships between cells (**Figure 4f**), as for ICICLE-seq, above (**Figure 3a,b**). However, these views do not leverage the strength of linked multimodal measurements. To begin to take advantage of our tri-modal measurements, we extended a method recently described by Y. Hao and S. Hao, *et al.* for paired weighted nearest-neighbors (WNN) analysis (Hao et al., 2020) to allow an arbitrary number of simultaneously measured modalities to contribute to the WNN network (**Methods**). After applying this three-way WNN, we could generate a UMAP embedding with contributions from all three simultaneously measured modalities (**Figure 4g**), which enhanced cell type separation. In this projection, we can view marker expression across all three modalities that is consistent with cell labels and cell surface phenotypes (e.g. CD8A accessibility, RNA expression, and cell surface quantification, **Figure 4h**), though we found that consistency across all modalities is not a universal characteristic of functional markers (**Figure 4 – figure supplement 2a**). In agreement with observations by Y. Hao and S. Hao, *et al.* on two-modality data, the weights contributed by different modalities to the WNN analyses vary by cell type (**Figure 4 – figure supplement 2b**), with T cell populations showing greater contribution from protein measurements to group cells in the WNN graph.

## Discussion

Multimodal data collection from permeabilized PBMCs will be of use to many researchers in the immunology field and beyond who seek to get the most high-quality data from precious clinical samples. We found that isotonic cell permeabilization allows the generation of scATAC-seq libraries with high quality as measured by FRIP and low nucleosomal content, suggesting that chromatin state in the nuclei is unperturbed (**Figure 1** and **Figure 2**). While a previous study utilized FACS followed by scATAC on individually sorted cells (Pi-ATAC; Xingqi Chen et al., 2018), the use of permeabilized cells enables simultaneous interrogation of chromatin accessibility state in the nucleus and the functional state of cells based on their cell surface proteins (ICICLE-seq, **Figure 3**) together with mRNA (TEA-seq, **Figure 4**) at unprecedented scale for the first time. Our methods utilize permeabilized cells to provide a direct link between high-quality scRNA-seq data and scATAC-seq data using truly paired methods akin to SNARE-seq (Chen et al., 2019) and SHARE-seq (Ma et al., 2020) by taking advantage of 10x Genomics Single Cell Multiome sequencing reagents as a platform for nucleic acid capture. While previous trimodal measurements have been performed using methods such as Patch-Seq (Cadwell et al., 2015) and scNMT-seq (Clark et al., 2018), the use of a droplet-based method allows multimodal measurements at a much higher scale by capturing thousands of single cells at once. Further development could allow simultaneous measurement of additional aspects of cell biology (e.g. CpG sequencing methods, scCUT&Tag (Bartosovic et al., 2020)). We anticipate that simultaneous, truly multimodal measurement across molecular compartments will be essential tools to expand our view of the full picture of immune cell state in health and disease, with many applications extending throughout the genomics research community.

## Author contributions

P.J.S., L.T.G., and E.S. designed the study. E.S. and C.L. performed scATAC-seq experiments. L.T.G. and R.G. performed scATAC-seq data processing. L.T.G. performed scATAC-seq analysis. E.S. designed and performed ICICLE-seq and TEA-seq experiments. E.S. and L.T.G. performed ICICLE-seq and TEA-seq analysis. R.G. and P.J.S. managed data repository submissions. J.R. performed FACS experiments. A.T.H. and J.R. performed flow cytometry. A.K.S., A.T.H., and J.R. designed flow cytometry panels and gating strategies. X.J.L. designed the multi-way WNN method and L.T.G. modified code for WNN implementation. T.F.B. provided oversight of the Allen Institute for Immunology. T.R.T. provided oversight of the AIFI Experimental Immunology group and edited the manuscript. L.T.G., E.S., and P.J.S. wrote the manuscript, with input from all coauthors.

## Acknowledgments

The authors thank Nina Kondza, Tanja Smith, Leila Shiraiwa, and Ernie Coffey for operational support, Morgan Weiss for assistance with preliminary experiments, the Allen Institute for Immunology Software Development team for computational support. The authors thank the Allen Institute founder, Paul G. Allen, for his vision, encouragement, and support.

## Supplement to

### Methods

#### Sample collection and preparation

##### Sample Collection and Processing

Biological specimens were purchased from BioIVT as cryopreserved PBMCs and Bloodworks NW as freshly drawn whole blood. All sample collections were conducted by BioIVT and Bloodworks NW under IRB-approved protocols, and all donors sign informed consent forms. See **Supplementary Table 7** for a list of sources and samples used for data displayed in each figure.

PBMCs sourced from BioIVT were isolated using either Ficoll-Paque or leukapheresis. Following isolation, PBMCs were subjected to RBC lysis, washing, and counting. PBMC aliquots were cryopreserved in Cryostor CS10 (StemCell Technologies, 07930) and stored in vapor phase liquid nitrogen.

For fresh blood samples from Bloodworks NW, PBMC processing occurred in-house. Blood tubes were pooled, gently swirled until fully mixed, about 30 times, and diluted with an equivalent volume of room temperature PBS (Thermo Fisher Scientific, 14190235). PBMCs were isolated using one or more Leucosep tubes (Greiner Bio-One, 227290) loaded with 15 mL of Ficoll Premium (GE Healthcare, 17-5442-03) to which a 3 mL cushion of PBS had been slowly added on top of the Leucosep barrier. Diluted whole blood (24-30mL) was slowly added to each tube and spun at 1000×g for 10 minutes at 20°C with no brake (Beckman Coulter Avanti J-15RIVD with JS4.750 swinging bucket, B99516). PBMCs were recovered from the Leucosep tube by quickly pouring all volume above the barrier into a sterile 50 mL conical tube (Corning, 352098). 15 mL cold PBS+0.2% BSA (Sigma, A9576; “PBS+BSA”) was added and the cells were pelleted at 400×g for 5-10 minutes at 4-10°C. The supernatant was quickly decanted, the pellet dispersed by flicking the tube, and the cells washed with 25-50 mL cold PBS+BSA. Cell pellets were combined as needed, the cells were pelleted as before, supernatant quickly decanted, and residual volume was carefully aspirated. PBMCs were resuspended in 1 mL cold PBS+BSA per 15 mL whole blood processed and counted with a ViCell (Beckman Coulter) using VersaLyse reagent (Beckman Coulter, A09777) or with a Cellometer Spectrum Cell Counter (Nexcelom) using ViaStain Acridine Orange/Propidium Iodide solution (Nexcelom, C52-0106-5). PBMCs were cryopreserved in Cryostor10 (StemCell Technologies, 07930) or 90% FBS (Thermo Fisher Scientific, 10438026) / 10% DMSO (Fisher Scientific, D12345) at 5×10^6^ cells/mL by slow freezing in a Coolcell LX (VWR, 75779-720) overnight in a −80°C freezer followed by transfer to liquid nitrogen.

##### Cell Thawing

Cryopreserved PBMCs were removed from liquid nitrogen storage and thawed in a 37°C water bath for 3-5 minutes until no ice was visible. Cells were diluted to 10 mL in 37°C AIM V medium (Gibco, 12055091) with the first 3 mL added dropwise. Cells were then washed once with 10 mL DPBS without calcium and magnesium (Corning, 21-031-CM) supplemented with 0.2% w/v BSA (Sigma-Aldrich, A2934). Cells were counted on a Cellometer Spectrum Cell Counter (Nexcelom) using ViaStain Acridine Orange/Propidium Iodide solution (Nexcelom, C52-0106-5) and stored on ice.

##### FACS Neutrophil Depletion

To remove dead cells, debris, and neutrophils, PBMC samples were sorted by fluorescence activated cell sorting (FACS) prior to nuclei isolation or cell permeabilization. Cells were incubated with Fixable Viability Stain 510 (BD, 564406) for 15 minutes at room temperature and washed with AIM V medium (Gibco, 12055091) plus 25mM HEPES before incubating with TruStain FcX (BioLegend, 422302) for 5 minutes on ice, followed by staining with anti-CD45 (BioLegend, 304038) and anti-CD15 (BD, 562371) antibodies for 20 minutes on ice. Cells were washed with AIM V medium plus 25mM HEPES and sorted on a BD FACSAria Fusion. A standard viable CD45+ cell gating scheme was employed; FSC-A v SSC-A (to exclude sub-cellular debris), two FSC-A doublet exclusion gates (FSC-W followed by FSC-H), dead cell exclusion gate (BV510 LIVE/DEAD negative) followed by CD45+ inclusion gate. Neutrophils (defined as SSC^high^, CD15+) were then excluded in the final sort gate (**Figure 1 - figure supplement 3**). An aliquot of each post-sort population was used to collect 50,000 events to assess post-sort purity.

##### Magnetic Bead Neutrophil Depletion

Bead-based neutrophil depletion was performed using a biotin conjugated monoclonal anti-CD15 antibody in combination with streptavidin coated magnetic beads. A high neutrophil content (approximately 1.1%) Ficoll isolated PBMC sample was processed to evaluate efficacy, and a low neutrophil leukapheresis isolated PBMC sample was processed to control for off-target effects. Briefly, 1×10^7^ PBMCs were resuspended in 100 μl of chilled DPBS without calcium and magnesium (Corning, 21-031-CM) supplemented with 0.2% w/v BSA (Sigma-Aldrich, A2934). 10 μl TruStain FcX (BioLegend 422302) was added to the cell suspension, mixed by pipette, and incubated on ice for 10 minutes. Anti-CD15 antibody (BioLegend, 301913) was added to the cell suspension, mixed by pipette, and incubated on ice for 30 minutes. Following antibody binding, 25 μl of Dynabeads MyOne Streptavidin T1 magnetic beads (Invitrogen, 65601) was added to the cell suspension, mixed by pipette, and incubated at room temperature for 5 minutes. The cell suspension was then diluted with 900 μl of room temperature DPBS+0.2% w/v BSA and placed on an EasySep magnet (Stemcell Technologies, 18103) for 3 minutes. The supernatant (approximately 1 ml) was transferred to a new tube and stored on ice until further processing.

Non-depleted and neutrophil depleted PBMCs from each sample were analyzed by flow cytometry using an 8-color panel to assess the effects of the bead-based depletion on major PBMC populations. For each sample and condition, 1×10^6^ cells were centrifuged (750×g for 5 minutes at 4°C) using a swinging bucket rotor (Beckman Coulter Avanti J-15RIVD with JS4.750 swinging bucket, B99516), the supernatant was removed using a vacuum aspirator pipette, and the cell pellet was resuspended in 100 μl of DPBS without calcium and magnesium (Corning, 21-031-CM) supplemented with 0.2% w/v BSA (Sigma-Aldrich, A2934). Cells were incubated with Fixable Viability Stain 510 (BD, 564406) and TruStain FcX (BioLegend, 422302) for 30 minutes on ice, and washed in chilled FACS buffer (DPBS, 0.2% w/v BSA, 0.1% sodium azide (VWR, BDH7465-2)). Cells were stained with a cocktail of antibodies (**Supplementary Table 1**) including 10 μl of Brilliant Stain Buffer Plus (BD, 566385) at a staining volume of 100 μl for 30 minutes on ice, then washed twice with chilled FACS buffer. Cells were passed through 35 μm Falcon Cell Strainers (Corning, 352235) and analyzed on a BD FACS Symphony flow cytometer. Gating analysis was performed using FlowJo cytometry software (Version 10.7).

A sequential gating scheme was used to identify viable singlet CD45+/CD15+/CD16+ neutrophils: 1, Time vs. SSC-A gate (to confirm that no abnormalities occurred in the fluidics), 2. FSC-A vs SSC-A (to exclude sub-cellular debris), 3. two FSC-A doublet exclusion gates (FSC-W followed by FSC-H), 3. dead cell exclusion gate (BV510 LIVE/DEAD negative) followed by 4. CD45+ inclusion gate.

Neutrophils were defined as either SSC-A^high^/CD15+ or CD15+/CD16+. The neutrophil population defined by SSC-A^high^/CD15+ was larger than that defined by CD15+/CD16+ due to the presence of some contaminating CD15^low^ monocytes. Therefore, we used the CD45+/CD15+/CD16+ gate for subsequent analysis including summary statistics. (**Figure 1 - figure supplement 4** and **Supplementary Table 2**).

##### Standard Nuclei Isolation

Isolation of nuclei suspensions was performed according to the Demonstrated Protocol: Nuclei Isolation for Single Cell ATAC Sequencing (10x Genomics, CG000169 Rev C). Briefly, 8×10^5^ to 1×10^6^ cells were added to a 1.5 mL low binding tube (Eppendorf, 022431021) and centrifuged (300×g for 5 minutes at 4°C) using a swinging bucket rotor (Beckman Coulter Avanti J-15RIVD with JS4.750 swinging bucket, B99516). The supernatant was removed using a vacuum aspirator pipette and the cell pellet was resuspended in 100 μl of chilled 10x Genomics Nuclei Isolation Buffer (10 mM Tris-HCl pH 7.4, 10 mM NaCl, 3 mM MgCl_2_, 0.1% Tween-20, 0.1 % NP-40 Substitute CAS 9016-45-9 (BioVision 2127-50), 0.01% Digitonin (MP Biomedicals 0215948082), 1% BSA) by pipette-mixing 10 times. Cells were incubated on ice for 3 minutes, followed by dilution with 1 mL of chilled 10x Wash Buffer (10 mM Tris-HCl pH 7.4, 10 mM NaCl, 3 mM MgCl_2_, 0.1% Tween-20 (BioRad 1610781), 1% BSA) by pipette-mixing 5 times. Nuclei were centrifuged (500×g for 3 minutes at 4°C) and the supernatant was slowly removed using a vacuum aspirator pipette. Nuclei were resuspended in chilled 1x Nuclei Buffer (10x Genomics, 2000207) to a target concentration of 3,000 - 6,000 nuclei per μl. Nuclei suspensions were passed through 35 μm Falcon Cell Strainers (Corning, 352235) and counted on a Cellometer Spectrum Cell Counter (Nexcelom) using ViaStain Acridine Orange/Propidium Iodide Staining Solution (Nexcelom, C52-0106-5).

##### Nuclei Isolation Optimization

In addition to 10x Nuclei Isolation Buffer (10xNIB), we tested an alternative Nuclei Isolation Buffer (ANIB) as described previously (Mulqueen et al., 2019: 10 mM Tris-HCl pH 7.4, 10 mM NaCl, 3 mM MgCl2, 0.1% Tween-20, 0.1 % IGEPAL CAS 9002-93-1 (Sigma, I8896), 1x Protease Inhibitor (Roche, 11836170001)). For each buffer, we generated a titration series of detergent concentrations relative to the concentrations described above but did not alter the concentration of other buffer ingredients: 1x, 0.5x, 0.25x, 0.1x for 10xNIB, and 1x and 0.1x for ANIB. The resulting nuclei were imaged using an EVOS M5000 Imaging System (Thermo Fisher Scientific, AMF5000) in transmitted light mode at 40x magnification to visually evaluate nuclear integrity (**Figure 1 - figure supplement 1**). The 1x 10xNIB, 0.25x 10xNIB, 0.1x 10xNIB, and 1x ANIB were used for 10X scATAC-seq (**Figure 1 - figure supplement 2**).

##### Cell Permeabilization

We prepared a 5% w/v digitonin stock by diluting powdered Digitonin (MP Biomedicals, 0215948082) with 100% DMSO (Fisher Scientific, D12345) and creating 20 μl aliquots which were stored at −20°C. To permeabilize, 1,000,000 cells were added to a 1.5 mL low binding tube (Eppendorf, 022431021) and centrifuged (400×g for 5 minutes at 4°C) using a swinging bucket rotor (Beckman Coulter Avanti J-15RIVD with JS4.750 swinging bucket, B99516). The supernatant was removed using a vacuum aspirator pipette and the cell pellet was resuspended in 100 μl of chilled isotonic Perm Buffer (20 mM Tris-HCl pH 7.4, 150 mM NaCl, 3 mM MgCl2, 0.01% Digitonin) by pipette-mixing ten times. Cells were incubated on ice for 5 minutes, after which they were diluted with 1 mL of isotonic Wash Buffer (20 mM Tris-HCl pH 7.4, 150 mM NaCl, 3 mM MgCl2) by pipette-mixing 5 times. Cells were centrifuged (400×g for 5 minutes at 4°C) using a swinging bucket rotor and the supernatant was slowly removed using a vacuum aspirator pipette. The cell pellet was resuspended in chilled TD1 buffer (Illumina, 15027866) by pipette-mixing to a target concentration of 2,300 - 10,000 cells per μl. Cells were passed through 35 μm Falcon Cell Strainers (Corning, 352235) and counted on a Cellometer Spectrum Cell Counter (Nexcelom) using ViaStain Acridine Orange/Propidium Iodide solution (Nexcelom, C52-0106-5). For optimization, we used varying final digitonin concentrations in the Perm Buffer: 0.01% w/v, 0.05% w/v, 0.1% w/v, and 0.2% w/v. The optimal concentration observed was 0.01% w/v.

#### snATAC-seq and scATAC-seq

##### sn/scATAC-seq Library Preparation

scATAC-seq libraries were prepared according to the Chromium Single Cell ATAC v1.1 Reagent Kits User Guide (CG000209 Rev B) with several modifications. 15,000 cells or nuclei were loaded into each tagmentation reaction. Nuclei were brought up to a volume of 5 μl in 1x Nuclei Buffer (10x Genomics, 2000207), mixed with 10 μl of a transposition master mix consisting of ATAC Buffer B (10x Genomics, 2000193) and ATAC Enzyme (Tn5 transposase; 10x Genomics, 2000123). Permeabilized cells were brought up to a volume of 9 μl in TD1 buffer (Illumina, 15027866) and mixed with 6 μl of Illumina TDE1 Tn5 transposase (Illumina, 15027916). Transposition was performed by incubating the prepared reactions on a C1000 Touch thermal cycler with 96–Deep Well Reaction Module (Bio-Rad, 1851197) at 37°C for 60 minutes, followed by a brief hold at 4°C. A Chromium NextGEM Chip H (10x Genomics, 2000180) was placed in a Chromium Next GEM Secondary Holder (10x Genomics, 3000332) and 50% Glycerol (Teknova, G1798) was dispensed into all unused wells. A master mix composed of Barcoding Reagent B (10x Genomics, 2000194), Reducing Agent B (10x Genomics, 2000087), and Barcoding Enzyme (10x Genomics, 2000125) was then added to each sample well, pipette-mixed, and loaded into row 1 of the chip. Chromium Single Cell ATAC Gel Beads v1.1 (10x Genomics, 2000210) were vortexed for 30 seconds and loaded into row 2 of the chip, along with Partitioning Oil (10x Genomics, 2000190) in row 3. A 10x Gasket (10x Genomics, 370017) was placed over the chip and attached to the Secondary Holder. The chip was loaded into a Chromium Single Cell Controller instrument (10x Genomics, 120270) for GEM generation. At the completion of the run, GEMs were collected and linear amplification was performed on a C1000 Touch thermal cycler with 96–Deep Well Reaction Module: 72°C for 5 min, 98°C for 30 sec, 12 cycles of: 98°C for 10 sec, 59°C for 30 sec and 72°C for 1 min.

GEMs were separated into a biphasic mixture through addition of Recovery Agent (10x Genomics, 220016), the aqueous phase was retained and removed of barcoding reagents using Dynabead MyOne SILANE (10x Genomics, 2000048) and SPRIselect reagent (Beckman Coulter, B23318) bead clean-ups. Sequencing libraries were constructed by amplifying the barcoded ATAC fragments in a sample indexing PCR consisting of SI-PCR Primer B (10x Genomics, 2000128), Amp Mix (10x Genomics, 2000047) and Chromium i7 Sample Index Plate N, Set A (10x Genomics, 3000262) as described in the 10x scATAC User Guide. Amplification was performed in a C1000 Touch thermal cycler with 96–Deep Well Reaction Module: 98°C for 45 sec, for 9 to 11 cycles of: 98°C for 20 sec, 67°C for 30 sec, 72°C for 20 sec, with a final extension of 72°C for 1 min. Final libraries were prepared using a dual-sided SPRIselect size-selection cleanup. SPRIselect beads were mixed with completed PCR reactions at a ratio of 0.4x bead:sample and incubated at room temperature to bind large DNA fragments. Reactions were incubated on a magnet, the supernatant was transferred and mixed with additional SPRIselect reagent to a final ratio of 1.2x bead:sample (ratio includes first SPRI addition) and incubated at room temperature to bind ATAC fragments. Reactions were incubated on a magnet, the supernatant containing unbound PCR primers and reagents was discarded, and DNA bound SPRI beads were washed twice with 80% v/v ethanol. SPRI beads were resuspended in Buffer EB (Qiagen, 1014609), incubated on a magnet, and the supernatant was transferred resulting in final, sequencing-ready libraries.

##### sn/scATAC-seq Sequencing

Final libraries were quantified using a Quant-iT PicoGreen dsDNA Assay Kit (Thermo Fisher Scientific, P7589) on a SpectraMax iD3 (Molecular Devices). Library quality and average fragment size was assessed using a Bioanalyzer (Agilent, G2939A) High Sensitivity DNA chip (Agilent, 5067-4626). Libraries were sequenced on the Illumina NovaSeq platform with the following read lengths: 51nt read 1, 8nt i7 index, 16nt i5 index, 51nt read 2.

##### sn/scATAC-seq Data Processing

Demultiplexing of raw base call files into FASTQ files was performed using 10x cellranger-atac mkfastq (10x Genomics v.1.1.0). To assess samples at an equal sequencing depth, FASTQ files were downsampled to a uniform total raw read count among compared samples: 2×10^8^ fragments for comparison of nuclei and cells across FACS conditions (**Figure 1 and 2**); 1.25×10^8^ fragments for optimization experiments (**Figure 1 - figure supplement 2**) due to lower available total read depth after sequencing. 10x cellranger-atac count was used to process sequencing reads by performing adapter trimming and sequence alignment to the GRCh38 (hg38) reference genome (refdata-cellranger-atac-GRCh38-1.1.0). The output files fragments.tsv.gz and singlecell.csv were utilized for downstream processing and quality control analysis.

To evaluate quality control metrics across all scATAC-seq datasets, we utilized bedtools (v2.29.1) and GNU parallel (Tange, 2011) v20161222 to generate overlap counts and feature count matrices for a panel of reference genomic regions: 518,766 peaks from a previous study of PBMCs by scATAC-seq (Lareau et al., 2019; supplementary file GSE123577_pbmc_peaks.bed.gz from GEO accession GSE123577) were converted from hg19 to hg38 coordinates using the UCSC liftOver tool (Hinrichs et al., 2006; kent source v402) and used to compute a standardized fraction of reads in peaks score for cells in each dataset (FRIP); 33,496 transcription start site regions (TSS ± 2kb) from Hg38 ENSEMBL release 93 (Yates et al., 2020) were filtered to select genes used in the 10x Genomics cellranger GRCh38 reference for scRNA-seq (refdata-cellranger-GRCh38-3.0.0) and used to compute the fraction of reads in TSS (FRITSS); and a set of 3,591,898 reference DNase hypersensitive sites from ENCODE (Meuleman et al., 2020; ENCODE File ID ENCFF503GCK) were used to assess distal regulatory element accessibility. In addition, we generated tiled window counts across the genome in 5k, 20k, 100k bins.

##### sn/scATAC-seq Quality Control

Custom R scripts were used to assess and filter preprocessed scATAC-seq data along a variety of quality metrics. Cells or nuclei with > 1,000 uniquely aligned fragments, FRIP > 0.2, FRITSS > 0.2, and fraction of fragments overlapping ENCODE reference regions > 0.5 were retained for downstream analysis. Cells or nuclei that passed these QC cutoffs were used to generate sparse count matrices and filtered fragments.tsv.gz files for downstream analysis.

To examine aggregate TSS accessibility, we selected fragments from fragments.tsv.gz that overlapped TSS regions describe above (TSS ± 2kb). For plotting, fragments were separated using cell barcodes (and fragment length in the case of **Figure 1 - figure supplement 5**) into separate groups. Fragment positions were converted to positions relative to TSS (sensitive to transcript strand orientation), and the number of fragments overlapping each position were calculated.

To examine CTCF motif accessibility, CTCF motif locations were obtained from genome-wide motif scans of non-redundant TF motifs (Vierstra, 2020; https://resources.altius.org/~jvierstra/projects/motif-clustering/releases/v1.0). Motifs were filtered to select CTCF motifs that overlapped ENCODE reference regions (Meuleman et al., 2020; ENCODE File ID ENCFF503GCK). Selected were ranked by their MOODS match, and the top 100,000 motifs were selected for analysis. Motif locations were expanded to a total of 4kb centered on the middle of each CTCF motif (using the resize function from GenomicRanges in R). Fragments from cells that passed QC filtering were converted to target site duplication (TSD) center positions (+5 bp from the 5’ end and −4 bp from the 3’ end of each fragment). All TSD centers that overlapped expanded CTCF motif regions were selected, and the number of TSD centers that overlapped each position relative to the CTCF motif were calculated (sensitive to CTCF motif strand orientation).

##### sn/scATAC-seq Dimensionality Reduction

For 2D projections of scATAC-seq data, we used binarized sparse matrices of 20kb window accessibility across the hg38 genome (excluding mitochondrial regions, chrM). Independently for each dataset, we selected features found in > 3% of cells/nuclei, weighted features using term frequency - inverse document frequency, log-transformed the resulting weights, and performed PCA using singular value decomposition to generate 50 reduced dimensions as described previously (Cusanovich et al., 2018; Hill, 2019). We then removed the first PC, which was strongly correlated with the number of available fragments and retained the remaining PCs up to PC 30. For display, we further reduced the dimensionality of selected PCs using UMAP(Becht et al., 2019; McInnes et al., 2018) (R package uwot, v0.1.8, parameters: scale = TRUE, min_dist = 0.2).

##### sn/scATAC-seq Cell Type Labeling

Labeling of scATAC-seq datasets was performed using the ArchR package (Granja et al., 2020) v0.9.4. In brief, filtered fragments.tsv.gz files after quality control were used to generate an ArchR GeneScore matrix and a tiled genome feature matrix for each dataset. Cells were grouped by performing iterative latent semantic indexing (LSI) on the tile matrix, followed by the shared nearest neighbor clustering approach implemented in Seurat (Stuart et al., 2019) v3.1.5. GeneScore data was then used to compare scATAC-seq clusters to a labeled reference scRNA-seq dataset consisting of 9,380 PBMCs generated by 10x Genomics, with labels provided by the Satija lab (https://www.dropbox.com/s/zn6khirjafoyyxl/pbmc_10k_v3.rds?dl=0) using ArchR’s implementation of the FindTransferAnchors method from Seurat. The best-scoring labels for each scATAC-seq cluster were used for downstream analysis and display (**Figure 2**), and label transfer scores for individual cells were used to compare label transfer between methods (**Figure 2c**).

##### sn/scATAC-seq Peak Analysis

After labeling cell types in each dataset, peaks for each cell type were generated using the ArchR functions addGroupCoverages and addReproduciblePeakSet. Within each dataset, peaks scores from each pair of cell types were compared using getMarkerFeatures performed in each direction separately by swapping foreground and background cell types. Differentially accessible peaks (DAPs) from each comparison were selected with filter string “FDR <= 0.05 & Log2FC >= log2(1.5)”.

To identify enriched TFBS motifs, CIS-BP motif annotations (Weirauch et al., 2014) were attached to each peak identified by ArchR using the addMotifAnnotations. Marker peaks for each cell type were identified using getMarkerFeatures without specifying foreground and background groups. Enriched motifs were identified using the ArchR function peakAnnoEnrichment (parameters: peakAnnotation = “Motif”, cutOff = “FDR <= 0.01 & Log2FC >= log2(1.5)”). Up to the top 10 enriched motifs for each cell type were plotted using the ArchR function plotEnrichHeatmap.

#### Cell Type Flow Cytometry

To assess cell type proportions, PBMCs were analyzed with a 25-color immunophenotyping flow cytometry panel. 1×10^6^ thawed PBMCs were centrifuged (750×g for 5 minutes at 4°C) using a swinging bucket rotor (Beckman Coulter Avanti J-15RIVD with JS4.750 swinging bucket, B99516), the supernatant was removed using a vacuum aspirator pipette, and the cell pellet was resuspended in DPBS without calcium and magnesium (Corning 21-031-CM). Cells were incubated with Fixable Viability Stain 510 (BD, 564406) and TruStain FcX (BioLegend, 422302) for 30 minutes at 4°C, then washed with chilled PBS+0.2% BSA (Sigma, A9576; “PBS+BSA”). Cells were stained with a cocktail of antibodies (**Supplementary Table 3**) at a staining volume of 100 μl for 30 minutes at 4°C, then washed with PBS+0.2% BSA. Fixation was performed by resuspending cells in 100 μl of 4% Paraformaldehyde (Electron Microscopy Sciences, 15713) and incubating for 15 minutes at 25°C, protected from light. Following fixation, cells were washed twice with PBS+0.2% BSA and resuspended in 100 μl PBS (without BSA). Stained cells were analyzed on a 5 laser Cytek Aurora spectral flow cytometer. Spectral unmixing was calculated with pre-recorded reference controls using Cytek SpectroFlo software (Version 2.0.2). Cell types were quantified by traditional bivariate gating analysis performed with FlowJo cytometry software (Version 10.7, **Figure 2 - figure supplement 1**).

#### ICICLE-seq

##### Tn5 Complexing

The assembly of Tn5 transposomes was performed as previously described (Mulqueen et al., 2019). DNA complexes containing mosaic-end sequences with either a poly-T or Nextera R2N 5’ overhang (Poly-T Top-L/MOSAIC_Bot, Tn5ME-s7_Top/MOSAIC_Bot) were created by annealing equimolar amounts of top and bottom oligos (**Figure 3 - figure supplement 1** and **Supplementary Table 5**) on a C1000 Touch thermal cycler with 96-Deep Well Reaction Module (Bio-Rad, 1851197) at 95°C for 5 minutes followed by 5°C decreases every 2 minutes until the temperature reached 20°C. Oligos were annealed at a concentration of 16 μM in 2x Dialysis Buffer (100 mM HEPES-KOH pH 7.5 (Teknova, 550000-016), 200 mM NaCl, 0.2 mM EDTA, 2 mM DTT (IBI Scientific, 21040), 0.2% Triton X-100 (Sigma-Aldrich, T8787), 20% Glycerol (Teknova, G1798)). Annealed complexes were mixed 1:1 for a final concentration of 8 nM. Tn5 transposase (Beta Lifescience, TN5-BL01) was supplemented with 5M NaCl at a final volume ratio of 1:8 NaCl to Tn5. The resulting NaCl/Tn5 mixture was mixed with the annealed complexes at a volume ratio of 1.2:1 ratio of DNA complexes to Tn5 and incubated at 25°C for 60 minutes to form final, reaction ready Tn5 complexes, which were stored at −20°C until use.

##### Antibody Staining

PBMCs were depleted of neutrophils, dead cells, and debris through FACS as described above. 2×10^6^ sorted PBMCs were centrifuged (400×g for 5 minutes at 4°C) using a swinging bucket rotor (Beckman Coulter Avanti J-15RIVD with JS4.750 swinging bucket, B99516), the supernatant was removed using a vacuum aspirator pipette, and the cell pellet was resuspended in 100 μl of DPBS without calcium and magnesium (Corning 21-031-CM) supplemented with 0.2% w/v BSA (Sigma-Aldrich A2934). 10 μl TruStain FcX (BioLegend, 422302) was added and cells were incubated on ice for 10 minutes. A panel of 46 barcoded oligo-conjugated antibodies (BioLegend TotalSeq-A) including a mouse IgG1ĸ isotype negative control (**Supplementary Table 6**) was added and incubated on ice for 30 minutes. Cells were washed three times in 4 mL of DPBS plus 2% BSA to remove unbound antibodies and used as input into cell permeabilization with 0.01% digitonin as described above.

##### ICICLE-seq Library Preparation

Transposition was performed by aliquoting 20,000 permeabilized cells in TD1 buffer (Illumina, 15027866), bringing the volume up to 9 μl in TD1 buffer, and mixing with 6 μl of Poly-T overhang Tn5 complexes. Reactions were incubated on a C1000 Touch thermal cycler with 96-Deep Well Reaction Module (Bio-Rad, 1851197) at 37°C for 120 minutes, followed by a brief hold at 4°C. Cell barcodes were then added to ATAC and antibody derived tags (ADTs) via GEM generation using 10x Genomics 3’ RNA beads and subsequent amplification. Briefly, a Chromium Next GEM Chip G (10x Genomics, 2000177) was placed in a Chromium Next GEM Secondary Holder (10x Genomics, 3000332) and 50% Glycerol (Teknova, G1798) was dispensed into all unused wells. A barcoding master mix was prepared which consisted of NEBNext Ultra II Q5 Master Mix (New England Biolabs, M0544), Reducing Agent B (10x Genomics, 2000087), F BC Primer (0.2 μM **Supplementary Table 5**), and ADT-Rev-AMP (0.2 μM **Supplementary Table 5**). The master mix was added to each sample well, pipette-mixed, and loaded into row 1 of the chip. Chromium Single Cell 3’ v3.1 Gel Beads (10x Genomics, 2000164) were vortexed for 30 seconds and loaded into row 2 of the chip, along with Partitioning Oil (10x Genomics, 2000190) in row 3. A 10x Gasket (10x Genomics, 370017) was placed over the chip and attached to the Secondary Holder. The chip was loaded into a Chromium Single Cell Controller instrument (10x Genomics, 120270) for GEM generation. At the completion of the run, GEMs were collected and amplification was performed on a C1000 Touch thermal cycler with 96-Deep Well Reaction Module: 72°C for 5 min, 98°C for 30 sec, 12 cycles of: 98°C for 10 sec, 42°C for 30 sec and 65°C for 30 sec, followed by a final extension of 65°C for 1 min.

GEMs were separated into a biphasic mixture through addition of Recovery Agent (10x Genomics, 220016), the aqueous phase was retained and removed of barcoding reagents using Dynabead MyOne SILANE (10x Genomics, 2000048) beads. Next, a dual-sided 0.6x/2.0x bead:sample SPRIselect reagent (Beckman Coulter, B23318) size-selection clean-up was performed to remove large DNA fragments and unused primers. Libraries were split into two reactions in a 3:1 ATAC:ADT ratio and amplified separately using different indexed P7 primers. ATAC fragments were amplified in a 100 μl reaction consisting of Buffer EB (Qiagen, 1014609), Amp Mix (10x Genomics, 2000047), SI-P5-22 primer (20 μM **Supplementary Table 5**), and Chromium i7 Multiplex Kit N Set A (10x Genomics, 3000262). ATAC PCR was performed in a C1000 Touch thermal cycler with 96-Deep Well Reaction Module: 98°C for 45 sec, 7 cycles of: 98°C for 20 sec, 54°C for 30 sec, 72°C for 20 sec, followed by a final extension of 72°C for 1 min. ADT fragments were amplified in a 100 μl reaction consisting of Buffer EB (Qiagen, 1014609), KAPA HiFi HotStart ReadyMix (KAPA Biosystems, KM2602), SI-P5-22 primer (10 μM **Supplementary Table 5**), and ADT i7 primer (10 μM **Supplementary Table 5**). ADT PCR was performed in a C1000 Touch thermal cycler with 96-Deep Well Reaction Module: 95°C for 3 min, 15 cycles of: 95°C for 20 sec, 60°C for 30 sec, 72°C for 20 sec, followed by a final extension of 72°C for 5 min. SPRIselect reagent cleanups were performed with a 1.2x bead:sample ratio for ADT libraries and a dual-sided size-selection of 0.4x/1.2x bead:sample ratio for ATAC libraries.

##### ICICLE-seq Sequencing

Final libraries were quantified using qPCR (KAPA Biosystems Library Quantification Kit for Illumina, KK4844) on a CFX96 Touch Real-Time PCR Detection System (Bio-Rad, 1855195). Library quality and average fragment size were assessed using a Bioanalyzer (Agilent, G2939A) High Sensitivity DNA chip (Agilent, 5067-4626). Libraries were sequenced on the Illumina NovaSeq platform with the following read lengths: 28 bp read 1 (Cell barcode and UMI), 8 bp i7 index, 100 bp read 2 (ATAC-seq sequence or ADT barcode). A Truseq read 1 primer (0.3 μM **Supplementary Table 5**) was included as a Custom Read 1 primer to mitigate the risk of off-target priming of the standard Illumina Nextera read 1 primer on the partial Nextera R1N sequence included in the mosaic end portion of the Poly-T Tn5 insertion.

##### ICICLE-seq Data Preprocessing

Demultiplexing of raw base call files into FASTQ files was performed using bcl2fastq2 (Illumina v2.20.0.422). Read 2 was trimmed of adapter sequences, low quality bases and reads, and polyA tailing using fastp (S. Chen et al., 2018) v0.21.0 (parameters: -- adapter_sequence=CTGTCTCTTATACACATCT --cut_tail --trim_poly_x) and the resulting read 2 sequences were aligned to the GRCh38 (hg38) reference genome (Illumina iGenomes, https://support.illumina.com/sequencing/sequencing_software/igenome.html) using Bowtie 2 (Langmead and Salzberg, 2012; v2.3.0, parameters: --local, --sensitive, --no-unal, --phred33). Aligned reads in SAM format were filtered by alignment score (greater than or equal to 30) then tagged with cell barcode and UMI sequence and quality scores using custom python code (python3 v3.7.3). Barcode sequences were compared against the 10x Genomics v3 3’ GEX barcode whitelist (3M-february-2018.txt.gz). Sequences not included in the whitelist were corrected to a valid whitelist barcode by allowing a single base mismatch (Hamming distance of 1). Sequences with more than one possible match were corrected at the position with the lowest sequencing quality score. Reads with barcodes that could not be corrected were excluded from further analysis. Filtered and tagged SAM files were converted to sorted, indexed BAM files using GATK (McKenna et al., 2010; Broad Institute v4.1.4.0). Genomic coordinates were converted to BED format using bedtools (Quinlan and Hall, 2010; v2.26.0). Custom python code was used to collapse aligned fragments into a list of fragments with unique cell barcode and genomic coordinate combinations. These fragments were then written as a fragments.tsv.gz file in the format: chr, start position, end position, cell barcode, UMI count, and strand (+/−).

##### ADT Data Preprocessing

Methods for ADT counting were developed in-house and were implemented as an optimized, highly efficient C program named BarCounter (available at https://github.com/AllenInstitute/Barcounter-release). BarCounter was used for single-cell ADT counting as follows: firstly, barcode sequences were compared against the 10x Genomics v3 3’ GEX barcode whitelist (3M-february-2018.txt.gz). Sequences not included in the whitelist were corrected to a valid whitelist barcode by allowing a single base mismatch (hamming distance of 1) at a low quality basecall (sequencing quality score < 20). Reads with barcodes that could not be corrected were excluded from further analysis. Next, ADT barcode sequences were compared against a CSV taglist containing ADT barcode / antibody associations. Antibody barcodes in the current TotalSeq-A catalog (BioLegend) have a Hamming distance from all other barcodes of at least 3. Therefore, a single base mismatch (Hamming distance of 1) was allowed. Reads containing ADT sequences that could not be assigned to an antibody in the taglist were excluded. Finally, UMI sequences that were unique within their assigned ADT for their assigned cell barcode were counted. Final ADT UMI counts were written by cell barcode to a CSV file for use in downstream analysis.

ADT features were filtered by comparison to the mouse IgG1ĸ isotype control, which should not bind to human cell surface proteins. The distribution of counts for each antibody was compared to the control using a Mann-Whitney test (R function wilcox.test with parameter alternative = “greater”). Any features for which the test returned a p-value > 1×10^−9^ were considered similar to the control and were removed from downstream analysis (**Supplementary Table 6**).

##### ICICLE-seq Analysis

Aligned ICICLE-seq chromatin accessibility fragments.tsv.gz files were preprocessed as for 10x scATAC-seq samples, above. QC filtering was performed as described, with modified cutoffs: > 500 uniquely aligned reads, FRIP > 0.65, FRITSS > 0.2, and fraction of fragments overlapping ENCODE reference regions > 0.5 were retained for downstream analysis. For 2D projections of the scATAC-seq data, we used binarized sparse matrices of 20kb window accessibility across the hg38 genome, selected features found in > 0.5% of cells, weighted features using LogTF-IDF, and performed PCA as described above. We then removed the first PC and retained the remaining PCs up to PC 20. UMAP was performed with adjusted parameters (scale = FALSE, min_dist = 0.2). To assign cell type labels, filtered fragments.tsv.gz files were used as input to ArchR. ArchR functions addIterativeLSI and addGeneIntegrationMatrix (parameters transferParams = list(dims = 1:10, k.weight = 20) and nGenes = 4000) were used to transfer labels from the scRNA-seq PBMC reference described above (scATAC Cell Type Labeling).

Count matrices for ADT data were scaled for each cell by dividing by the thousands of total ADT UMIs per cell, then transformed using Log10(scaled count + 1). Normalized features were used for PCA using the R function prcomp with default parameters. Filtered and normalized features were used as direct input to UMAP (R package uwot with parameter min_dist = 0.2) with the first 2 PCs from PCA used as initial coordinates to aid reproducibility of UMAP projection. Cells were clustered using a Jaccard-Louvain method (parameters k = 15, radius = 1) using UMAP coordinates. Clusters with high signal from the mouse IgG1κ isotype control antibody were removed from subsequent analysis (1 cluster, n = 32 cells). The remaining clusters were manually labeled by examination of cell type marker enrichment. The R package scratch.vis (Tasic et al., 2018; https://github.com/alleninstitute/scrattch.vis) was used to generate the cluster median heatmap plot and a river/alluvial plot comparing the cell type labels obtained from ATAC-seq and ADT-based analyses.

To compare peaks between cell types, all filtered fragments for cells in each cell type were aggregated and used as input to the MACS2 peak caller (Zhang et al., 2008; parameters -f BED, -g hs, --no-model). The top 2,500 peaks from each cell type were selected for comparison (except for Plasmablasts, for which all 592 peaks were used). A master set of peaks across all types was constructed by combining all narrowPeak results files and combining the outer coordinates of overlapping peaks (GenomicRanges function reduce). A binary matrix of peak overlaps for each cell type was generated and used to construct the peak comparison figure inspired by UpSet plots (Lex et al., 2014).

#### TEA-seq

##### TEA-seq Library Preparation

Consistent with ICICLE-seq, cryopreserved PBMCs were thawed, depleted of neutrophils, dead cells, and debris through FACS, and permeabilized using 0.01% digitonin as described above. ATAC and Gene Expression libraries were prepared according to the Chromium Next GEM Single Cell Multiome ATAC + Gene Expression User Guide (CG000338 Rev A) with several modifications. Transposition was performed by diluting 15,400 permeabilized cells to a final volume of 5 μl in isotonic Tagmentation Buffer (20 mM Tris-HCl pH 7.4, 150 mM NaCl, 3 mM MgCl_2_, RNase Inhibitor 1U/μl (Lucigen NxGen, F83923-1)). Cells were mixed with 10 μl of a transposition master mix consisting of 7 μl ATAC Buffer B (10x Genomics, 2000193) and 3 μl ATAC Enzyme B (Tn5 transposase; 10x Genomics, 2000265) per reaction. Transposition was performed by incubating the prepared reactions on a C1000 Touch thermal cycler with 96–Deep Well Reaction Module (Bio-Rad, 1851197) at 37°C for 60 minutes, followed by a brief hold at 4°C. A Chromium NextGEM Chip J (10x Genomics, 2000264) was placed in a Chromium Next GEM Secondary Holder (10x Genomics, 3000332) and 50% Glycerol (Teknova, G1798) was dispensed into all unused wells. A master mix composed of Barcoding Reagent Mix (10x Genomics, 2000267), Reducing Agent B (10x Genomics, 2000087), Template Switch Oligo (10x Genomics, 3000228), and Barcoding Enzyme Mix (10x Genomics, 2000266) was then added to each sample well, pipette-mixed, and loaded into row 1 of the chip. Chromium Single Cell Multiome Gel Beads v1.1 (10x Genomics, 2000261) were vortexed for 30 seconds and loaded into row 2 of the chip, along with Partitioning Oil (10x Genomics, 2000190) in row 3. A 10x Gasket (10x Genomics, 3000072) was placed over the chip and attached to the Secondary Holder. The chip was loaded into a Chromium Single Cell Controller instrument (10x Genomics, 120270) for GEM generation. At the completion of the run, GEMs were collected, and reverse transcription and barcoding were performed by incubating GEMs on a C1000 Touch thermal cycler with 96–Deep Well Reaction Module at 37°C for 45 min, 25°C for 30 min, followed by a brief hold at 4°C. Upon completion of the GEM incubation 5 μl of Quenching Agent (10x Genomics, 2000269) was immediately added and mixed with the reaction solution.

GEMs were separated into a biphasic mixture through addition of Recovery Agent (10x Genomics, 220016), the aqueous phase was retained and removed of barcoding reagents using Dynabead MyOne SILANE (10x Genomics, 2000048) and 2.0x bead:sample ratio SPRIselect reagent (Beckman Coulter, B23318) cleanups, incubating for 10 minutes after bead each addition step.

Barcoded ATAC and cDNA fragments were amplified using a PCR reaction consisting of 45 μl of template, 50 μl of Pre-Amp Mix (10x Genomics, 2000270), 5 μl of Pre-Amp Primers (10x Genomics, 2000271), and 1 μl of ADT-Rev-AMP (0.2 μM **Supplementary Table 5**). Amplification was performed in a C1000 Touch thermal cycler with 96–Deep Well Reaction Module: 72°C for 5 min, 98°C for 3 min, for 7 cycles of: 98°C for 20 sec, 63°C for 30 sec, 72°C for 1 min, with a final extension of 72°C for 1 min. Amplified fragments were purified using a 2.0x bead:sample ratio SPRIselect reagent (Beckman Coulter, B23318) bead clean-up, incubating for 10 minutes after bead addition.

Sequencing ready ATAC libraries were constructed by amplifying barcoded ATAC fragments in a sample indexing PCR consisting of SI-PCR Primer B (10x Genomics, 2000128), Amp Mix (10x Genomics, 2000047) and Sample Index Plate N, Set A (10x Genomics, 3000427) as described in the 10x Multiome User Guide. Amplification was performed in a C1000 Touch thermal cycler with 96–Deep Well Reaction Module: 98°C for 45 sec, for 9 cycles of: 98°C for 20 sec, 67°C for 30 sec, 72°C for 20 sec, with a final extension of 72°C for 1 min. Final ATAC libraries were prepared using a dual-sided 0.6x/1.6x bead:sample SPRIselect reagent (Beckman Coulter, B23318) size-selection clean-up.

cDNA fragments were amplified using a PCR reaction consisting of 35 μl of template, 50 μl of Amp Mix (10x Genomics, 2000047), 15 μl of cDNA Primers (10x Genomics, 2000089), and 1 μl of ADT-Rev-AMP (2 μM **Supplementary Table 5**). Amplification was performed in a C1000 Touch thermal cycler with 96–Deep Well Reaction Module: 98°C for 3 min, for 8 cycles of: 98°C for 15 sec, 63°C for 20 sec, 72°C for 1 min, with a final extension of 72°C for 1 min. cDNA and ADT fragments were separated using a dual-sided SPRIselect reagent (Beckman Coulter, B23318) size-selection clean-up. Large cDNA fragments were retained in an initial 0.6x bead:sample SPRIselect incubation, reactions were incubated on a magnet, and small unbound fragments containing ADTs were transferred to new wells where they were subjected to an additional 2.0x bead:sample SPRIselect cleanup.

Consistent with ICICLE-seq, ADT fragments were amplified in a 15 cycle indexing PCR. Final ADT libraries were prepared using a dual-sided 1.6x bead:sample SPRIselect reagent (Beckman Coulter, B23318) clean-up.

cDNA Fragmentation, End-Repair, and A-tailing were performed in a joint reaction containing 25% of cDNA sample as described in the 10x Multiome User Guide. Briefly, cDNA was diluted in Buffer EB (Qiagen, 1014609), and combined with master mix of Fragmentation Buffer (10x Genomics, 2000091) and Fragmentation Enzyme (10x Genomics, 2000090). The reaction was incubated in a C1000 Touch thermal cycler with 96–Deep Well Reaction Module: 4°C start, 32°C for 5 min, 65°C for 30 min, followed by a brief hold at 4°C. Reactions were purified using a dual-sided 0.6x/0.8x bead:sample SPRIselect reagent (Beckman Coulter, B23318) size-selection clean-up. Ligation was performed by mixing each sample with a master mix consisting of Ligation Buffer (10x Genomics, 2000092), DNA Ligase (10x Genomics, 220110), and Adapter Oligos (10x Genomics, 2000094), and incubating in a C1000 Touch thermal cycler with 96–Deep Well Reaction Module for 15 minutes at 20°C, followed by a brief hold at 4°C. Ligation reactions were purified using a 0.8x bead:sample SPRIselect reagent (Beckman Coulter, B23318) clean-up.

Sequencing ready Gene Expression libraries were constructed by amplifying cDNA fragments in a sample indexing PCR consisting of Amp Mix (10x Genomics, 2000047) and Dual Index TT Set A (10x Genomics, 3000431) as described in the 10x Multiome User Guide. Amplification was performed in a C1000 Touch thermal cycler with 96–Deep Well Reaction Module: 98°C for 45 sec, for 14 cycles of: 98°C for 20 sec, 54°C for 30 sec, 72°C for 20 sec, with a final extension of 72°C for 1 min. Final Gene Expression libraries were prepared using a dual-sided 0.6x/0.8x bead:sample SPRIselect reagent (Beckman Coulter, B23318) size-selection clean-up.

##### TEA-seq Sequencing

Final libraries were quantified using qPCR (KAPA Biosystems Library Quantification Kit for Illumina, KK4844) on a CFX96 Touch Real-Time PCR Detection System (Bio-Rad, 1855195). Library quality and average fragment size were assessed using a Bioanalyzer (Agilent, G2939A) High Sensitivity DNA chip (Agilent, 5067-4626). Libraries were sequenced on the Illumina NovaSeq platform with the following read lengths: 50 bp read 1, 10 bp i7 index, 16 bp i5 index, 90 bp read 2.

##### TEA-seq Data Processing

Demultiplexing of raw base call files into FASTQ files was performed using bcl2fastq2 (Illumina v2.20.0.422, parameters: --create-fastq-for-index-reads, --minimum-trimmed-read-length=8, --mask-short-adapter-reads=8, --ignore-missing-positions, -ignore-missing-filter, --ignore-missing-bcls, -r 24 -w 24 -p 80). The --use-bases-mask option was used to adjust the read lengths for each library type as follows: Y28n*,I10,I10n*,Y90n* for Gene Expression, Y50n*,I8n*,Y16,Y50n* for ATAC, and Y28n*,I8n*,n*,Y90n* for ADT.

##### TEA-seq Analysis

RNA and ATAC libraries were aligned using cellranger-arc software (v1.0.0, 10x Genomics) against 10x genomics reference refdata-cellranger-arc-GRCh38-2020-A using default parameters. ADT barcode counting was performed using BarCounter as described above for ICICLE-seq. After alignment, ATAC-seq data was used as input for scATAC-seq Quality Control analysis and filtering, as described above. 10x droplet barcodes that were called cells by the cellranger-arc software, passed ATAC-seq QC criteria (FRIP > 0.2, FRITSS > 0.2, and fraction of fragments overlapping ENCODE reference regions > 0.5), and had > 750 genes detected were used for downstream analysis.

Selected scATAC-seq barcodes were further processed using ArchR (v1.0.0) for Iterative LSI, Clustering, Group Coverage computation, Reproducible Peak Set annotation, and sparse Peak Matrix generation using default settings according to the ArchR documentation (https://archrproject.com).

Next, the scRNA-seq count matrix generated by cellranger-arc, the Peak Matrix (for analysis) and Gene Score Matrix (for visualization) generated using ArchR, and the ADT count matrix were filtered and combined as assays in a Seurat object (Seurat v4.0.0-beta), and dimensionality reduction was carried out using parameters specific to each assay as described below.

For scRNA-seq, the matrix was normalized (NormalizeData function, default settings), variable features were identified (FindVariableFeatures function, defaults), the data was scaled (ScaleData function, defaults), PCA was performed (RunPCA function, defaults), and a UMAP projection was generated (RunUMAP, dims = 1:30). Cell type labels were applied using Seurat label transfer using the scRNA-seq data, with the same Seurat reference dataset used for scATAC-seq above, by identifying transfer anchors (FindTransferAnchors function, reduction = “cca”), and transfering labels (TransferData function, weight.reduction = “cca”).

For ADT counts, all antibody features were selected as variable features, the data was normalized (NormalizeData function, scale.factor = 1000) and scaled (ScaleData function, defaults) before PCA (RunPCA function, npcs = 20) and UMAP projection (RunUMAP function, dims = 1:12).

For scATAC-seq data, we utilized the functions provided by the Signac package (v1.0.0) as follows: the binarized Peak Matrix was weighted (RunTFIDF function, defaults), top features were identified (RunTopFeatures, min.cutoff = ‘q75’), SVD was performed (RunSVD function, defaults), and SVD dimensions were used as input to UMAP projection (RunUMAP function, dims = 2:30).

##### Multi-modal Weighted Nearest Neighbors Analysis

In order to perform weighted nearest neighbors (WNN) analysis with input from all 3 assays above, we extended the functions provided by Seurat v4.0.0-beta (Hao et al., 2020), which utilize only two assays, to allow an arbitrary number of reduced dimension sets to be used as input. To do so, we performed within-modality and cross-modality predictions and computed cross-modal affinities as described by Y. Hao and S. Hao, *et al.* We then generalized the computation of cell-specific modality affinity ratio, *S*, for each cell i, in each modality, *α*, using a small value for *ε* (10^−4^) as suggested by Y. Hao and S. Hao, *et al.*

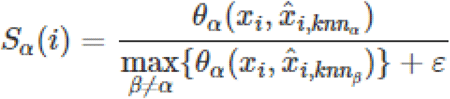

Here, *x*_*i*_ and 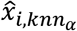 represent a generic form of the actual and predicted values of the low-dimensionality representation of each cell in each modality, as described for RNA and Protein in Y. Hao and S. Hao, *et al.* (e.g. *r*_*i*_ and 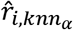 for scRNA), and *β* is used to represent all modalities that are not the modality under consideration (e.g. ADT and scATAC-seq when *α* is RNA). The formula for 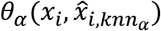 and 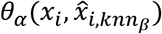 are given in Y. Hao and S. Hao, *et al.*

With these multiple *S*_*α*_(*i*) values, we then compute the cell-specific modality weights, *w*_*α*_(*i*) as:

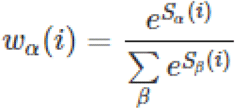

To generate the weighted pairwise cell similarities, we now sum across all modalities instead of only RNA and Protein:

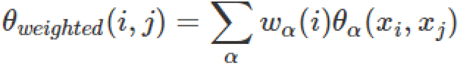

These multi-modal weighted similarities were then used to construct the WNN graph and perform WNN-based UMAP projection as described by Y. Hao and S. Hao, *et al.* The modified version of Seurat v4.0.0-beta used for this analysis is available at https://github.com/aifimmunology/seurat.

#### Data analysis and visualization software

Post-processing analysis of summary statistics and visualization of snATAC-seq, scATAC-seq, and ICICLE-seq was performed using R v.3.6.3 and greater (R Core Team) in the Rstudio IDE (RStudio Team, 2020; Integrated Development Environment for R) or using the Rstudio Server Open Source Edition as well as the following packages: for scATAC-seq specific analyses and comparisons to scRNA-seq data, ArchR (Granja et al., 2020) and Seurat (Stuart et al., 2019); for general data analysis and manipulation, data.table (Dowle and Srinivasan, 2019), dplyr, Matrix (Bates and Maechler, 2018), matrixStats (Bengtsson, 2018), purrr (Henry and Wickham, 2019), and reshape2 (Wickham, 2007); for data visualization, ggplot2 (Wickham, 2016), and cowplot (Wilke, 2018); for dimensionality reduction and clustering, igraph (Csardi and Nepusz, 2006), RANN, and the R uwot implementation (Melville, 2020) of UMAP (Becht et al., 2019; McInnes et al., 2018); for manipulation of genomic region data, bedtools2 (Quinlan and Hall, 2010) and GenomicRanges (Lawrence et al., 2013).

### Figure supplements

**Figure 1 – figure supplement 1.**
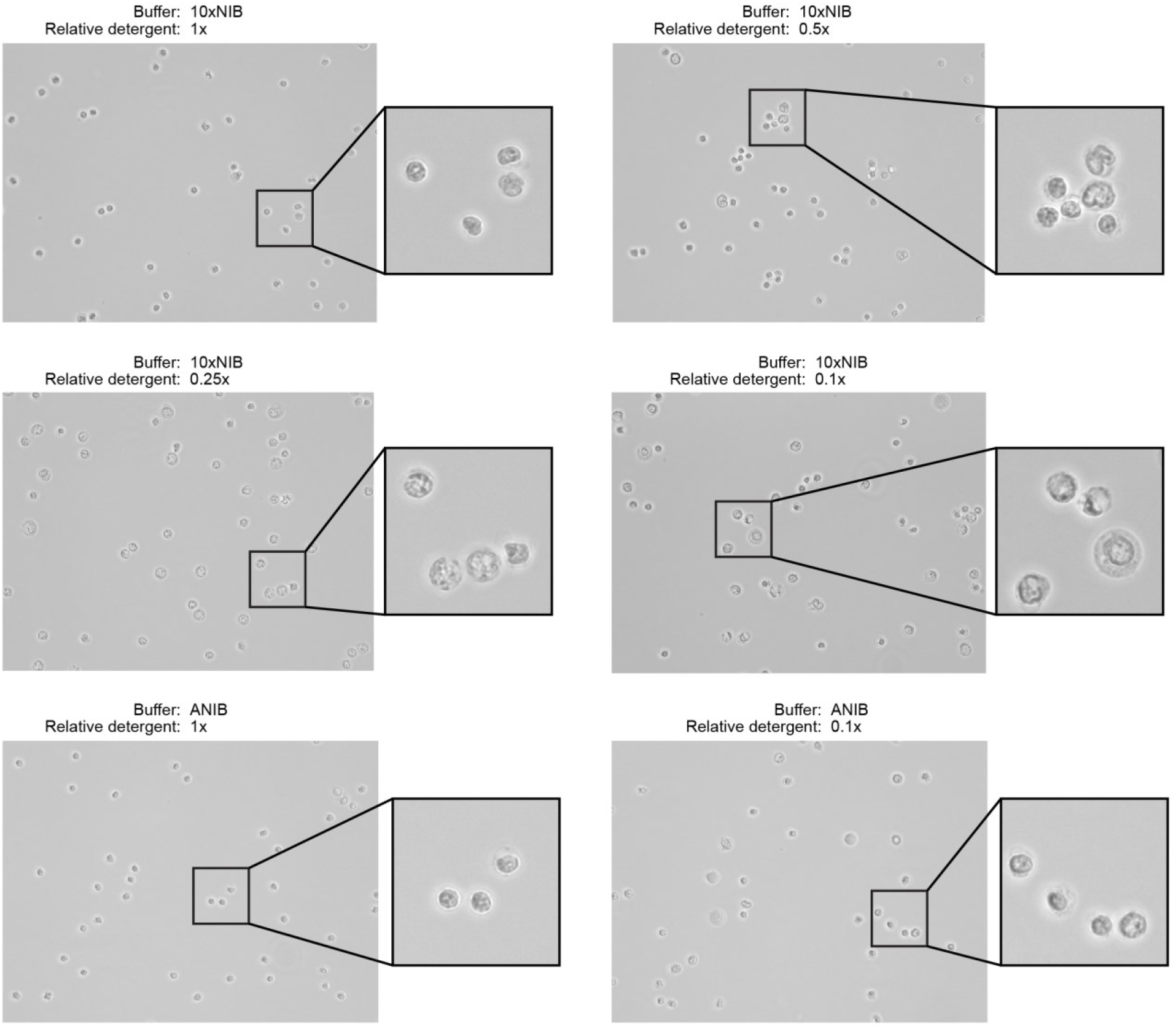
Nuclei were isolated from PBMCs using two buffer compositions and varying detergent concentrations. The two buffers, 10xNIB and ANIB, are defined in **Methods**, and the concentration of the detergent components of the buffers was varied for each sample. After isolation, nuclei were imaged using an EVOS M5000 Imaging System in transmitted light mode at 40x magnification to assess completeness of isolation and the condition of nuclei. Ideally, nuclei will not retain cell membranes and will have round, well-defined edges with minimal blebbing.

**Figure 1 – figure supplement 2.**
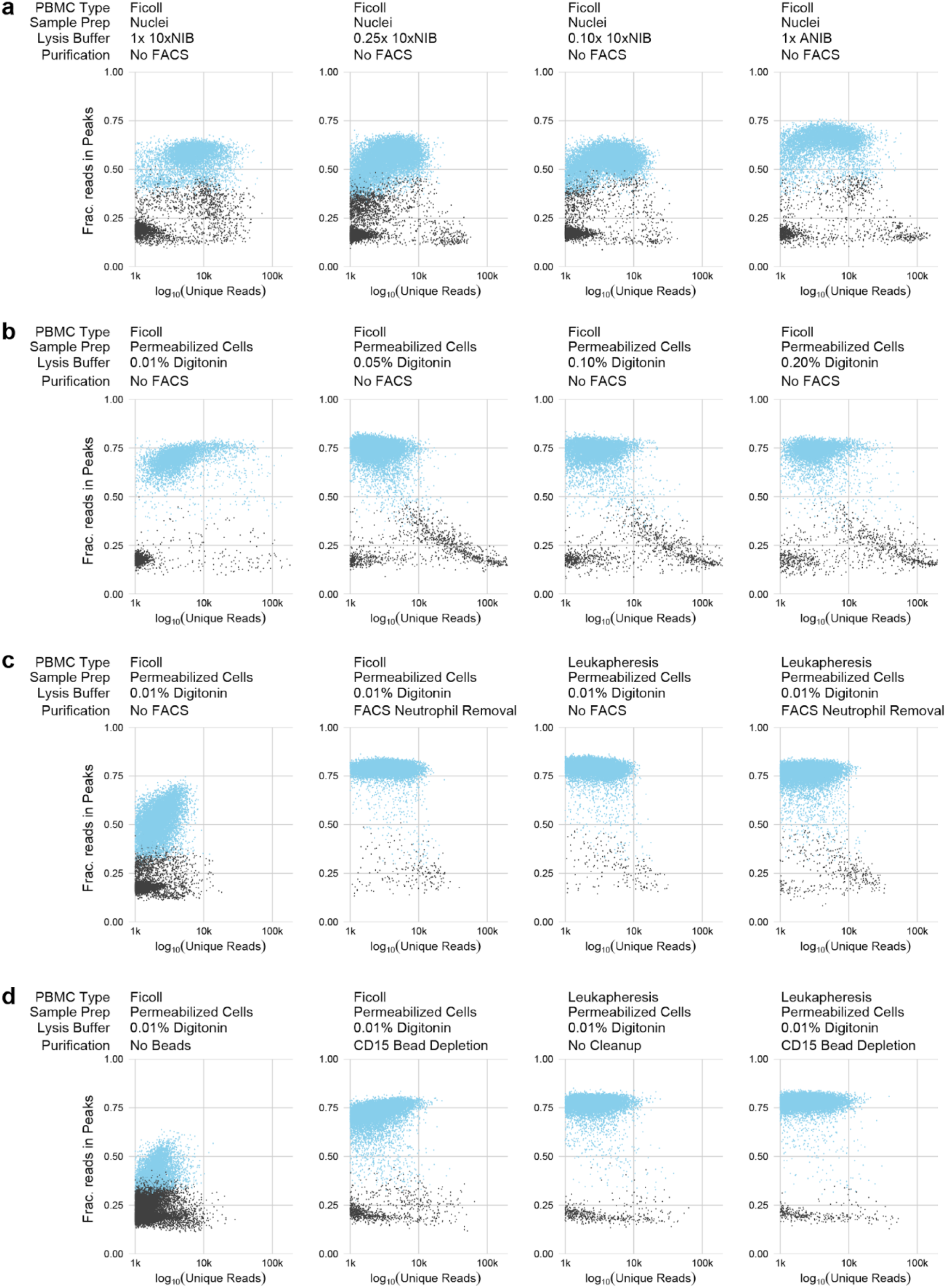
Quality control plots for snATAC-seq and scATAC-sec experimental conditions. Each plot compares the quantity of data obtained from each cell (x-axis, log-scaled unique fragment counts) against signal-to-noise as measured by fraction of reads in peaks (y-axis, FRIP). Each point represents a single 10x cell barcode. Blue points are barcodes that pass QC cutoffs (> 1,000 unique fragments, FRIP > 0.2, FRITSS > 0.2, and fraction of reads in ENCODE index peaks > 0.5); Gray points are barcodes that fail QC. **a**, Comparisons of nuclei isolation buffer compositions. Components of 10xNIB and ANIB buffers are defined in **Methods**. **b**, Comparisons of cell permeabilization conditions by varying the concentration of digitonin. Additional buffer components are defined in **Methods**. **c**, Comparisons of neutrophil removal by FACS on Ficoll-purified PBMCs (left 2 panels) and leukapheresis-purified PBMCs (right 2 panels). Neutrophil removal was performed using the gating scheme shown in **Figure 1 - figure supplement 3**. **d**, Comparisons of neutrophil removal by anti-CD15 magnetic bead depletion on Ficoll-purified PBMCs (left 2 panels) and leukapheresis-purified PBMCs (right 2 panels).

**Figure 1 – figure supplement 3.**
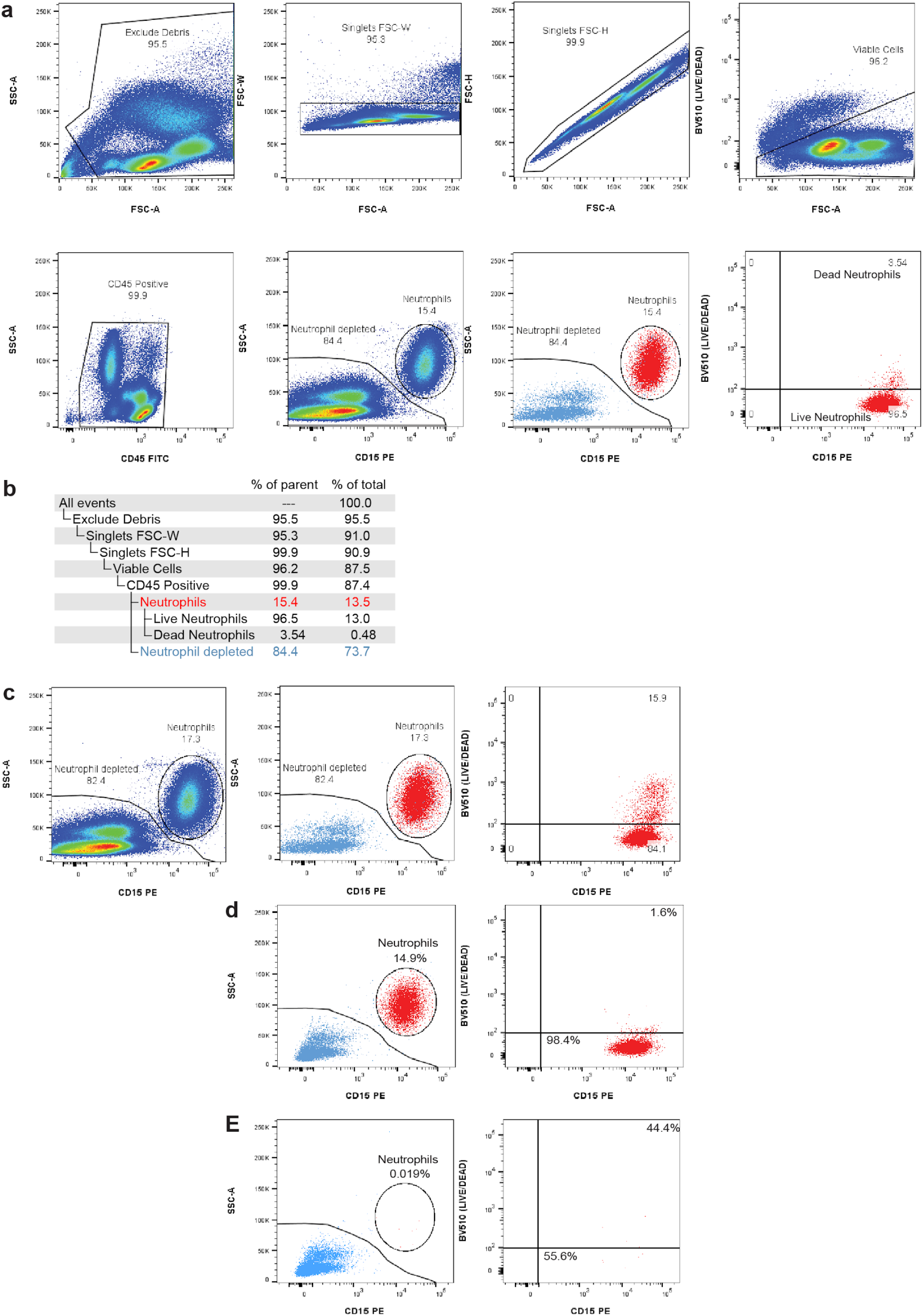
**a**, Gating strategy for FACS removal of debris, doublets, dead cells, and neutrophils. **b**, Gating hierarchy for FACS neutrophil depletion shown in **a**, with demonstration data used to provide percentages of parent gates and total cells. **c**, Pre-sort collection sequential gating shows the live/dead proportion of total neutrophils. During FACS, dead neutrophils are removed using the live/dead gate and CD15/SSC gate leading to an underestimation of the total neutrophil content in the sample. **d**, Post-sort analysis of neutrophil content after debris, doublet, and dead cell removal. Left panel displays the percentage of viable singlets in the neutrophil gate (CD45+ gate in **b**). The right panel displays the percentage neutrophils that are live (bottom-right) and dead (top-right). **e**, Post-sort analysis of neutrophil content and viability after neutrophil exclusion sort (Neutrophil depleted gate in **b**).

**Figure 1 – figure supplement 4.**
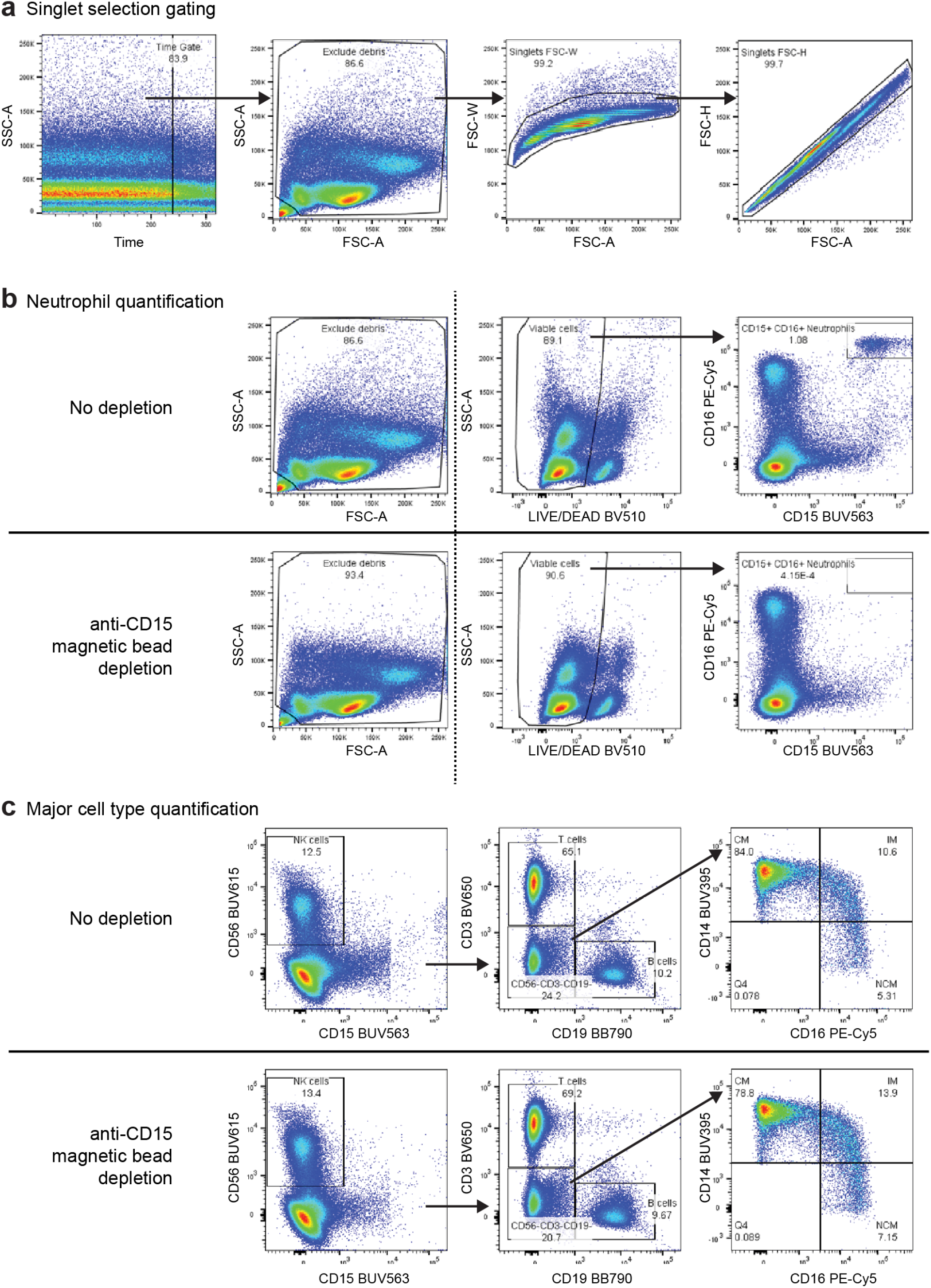
Gating strategy for the 8-color flow cytometry panel used to evaluate anti-CD15 bead-based neutrophil removal. **a**, Preparatory gating used to select high-quality events (plot 1), exclude debris (plot 2), and select singlet events (plots 3 and 4) **b**, Comparison of select gated populations of the PBMC sample before (top) and after depletion (bottom). The dashed line indicates that events from the left panel were not the parent gates for the two plots to the right, which were filtered first by the singlet gates described in **a**. Dead cells and debris were slightly reduced following depletion (Left-most plots). CD15+/CD16+ neutrophils were depleted from 1.080% to < 0.001% following depletion (right-most plots). **c**, Gating strategy for identification of major cells types in the PBMC sample before (top) and after depletion (bottom). Following time, debris, singlet, live/dead, and CD45+ gating, NK cells were defined as CD15-/CD56+, T cells were defined as CD3+/CD19-, and B cells were defined as CD3-/CD19+. Cells in the CD3-/CD19-/CD56-gate were used to identify monocyte subsets. Classical monocytes were defined as CD14+/CD16-, intermediate monocytes were defined as CD14+/CD16+, and non-classical monocytes were defined as CD14-/CD16+. We observed modest depletion of some classical monocytes (CM, from 84.0% to 78.8% of monocytes) with corresponding increases in the proportion of Intermediate (IM) and Non-classical (NCM) monocytes after bead depletion.

**Figure 1 – figure supplement 5.**
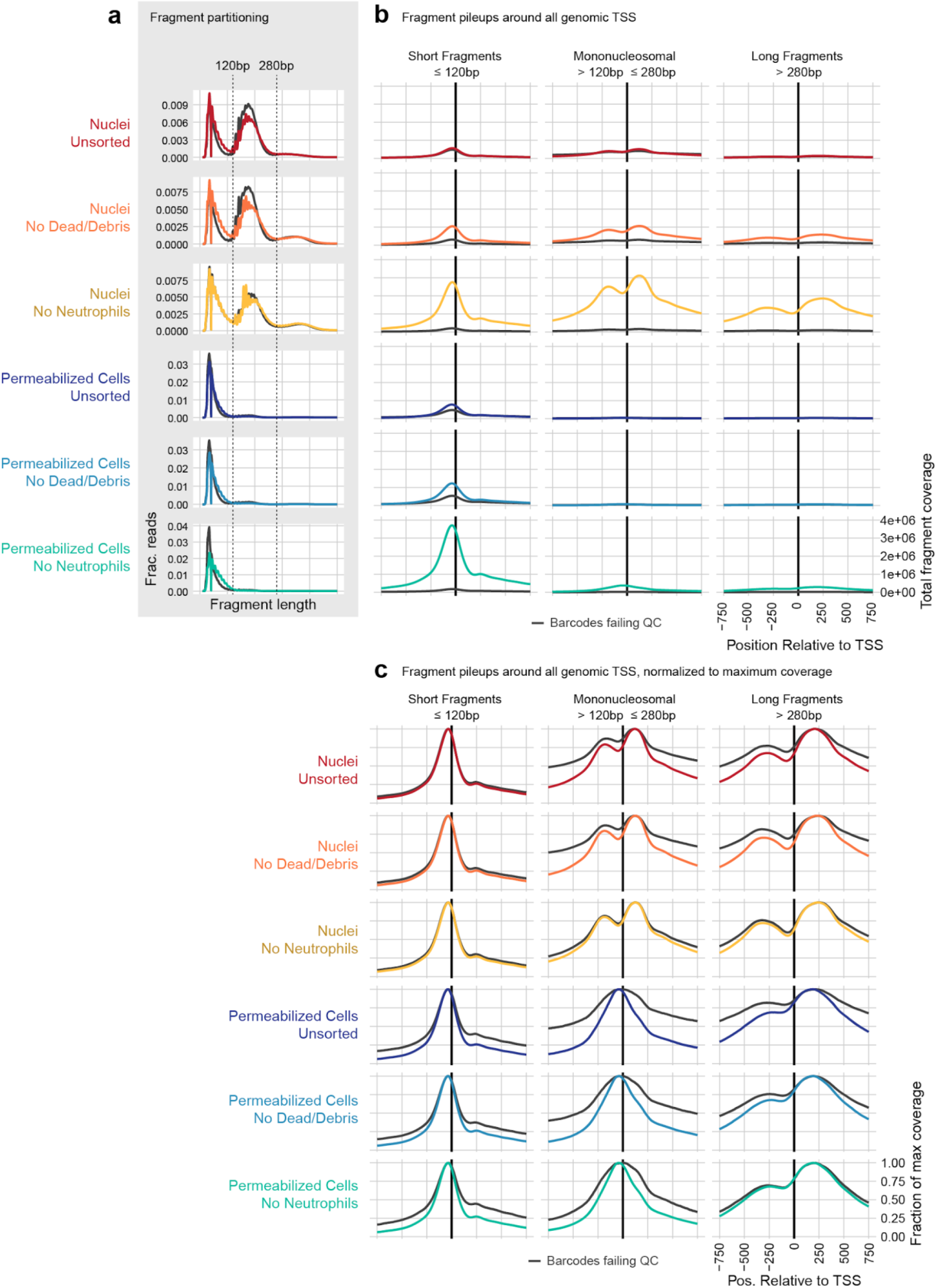
**a**, To compare the contribution of nucleosomal fragments to signal near genomic transcription start sites, we partitioned unique, aligned fragments into 3 categories: Short subnucleosomal fragments (≤120bp), mononucleosomal fragments (> 120bp and ≤ 280 bp), and longer fragments including di- and tri-nucleosomal fragments (> 280 bp). The distribution of fragment lengths for each sample is displayed as a line plot, where colored lines show the distribution of cell barcode fragments, and gray lines show the distribution of fragments from non-cell barcodes. **b**, All fragments that overlapped a window around all genomic TSS ± 2,000bp were assembled, and the overlaps of fragments in each of the 3 size categories defined in (a) are plotted relative to the TSS ± 750bp. Axis labels shown on the bottom-right panel apply to all plots in (b). Nucleosomal fragments flanking the TSS are clearly visible for the Nuclei: No Neutrophils sample (third row, gold). These are less frequent in all Permeabilized Cell samples (rows 4-6, blue colors). **c**, as for (b), but each line has been scaled by dividing by the maximum coverage value for that line to make the shape of each distribution visible. Here, we can see nucleosomal positioning in each of the Nuclei samples (top 3 rows, warm colors), and a lack of mononucleosmal positioning in Permeabilized Cell samples (bottom 3 rows, blue colors). However, longer flanking fragments (third column) have a similar qualitative shape in both Nuclei and Permeabilized cells.

**Figure 2 – figure supplement 1.**
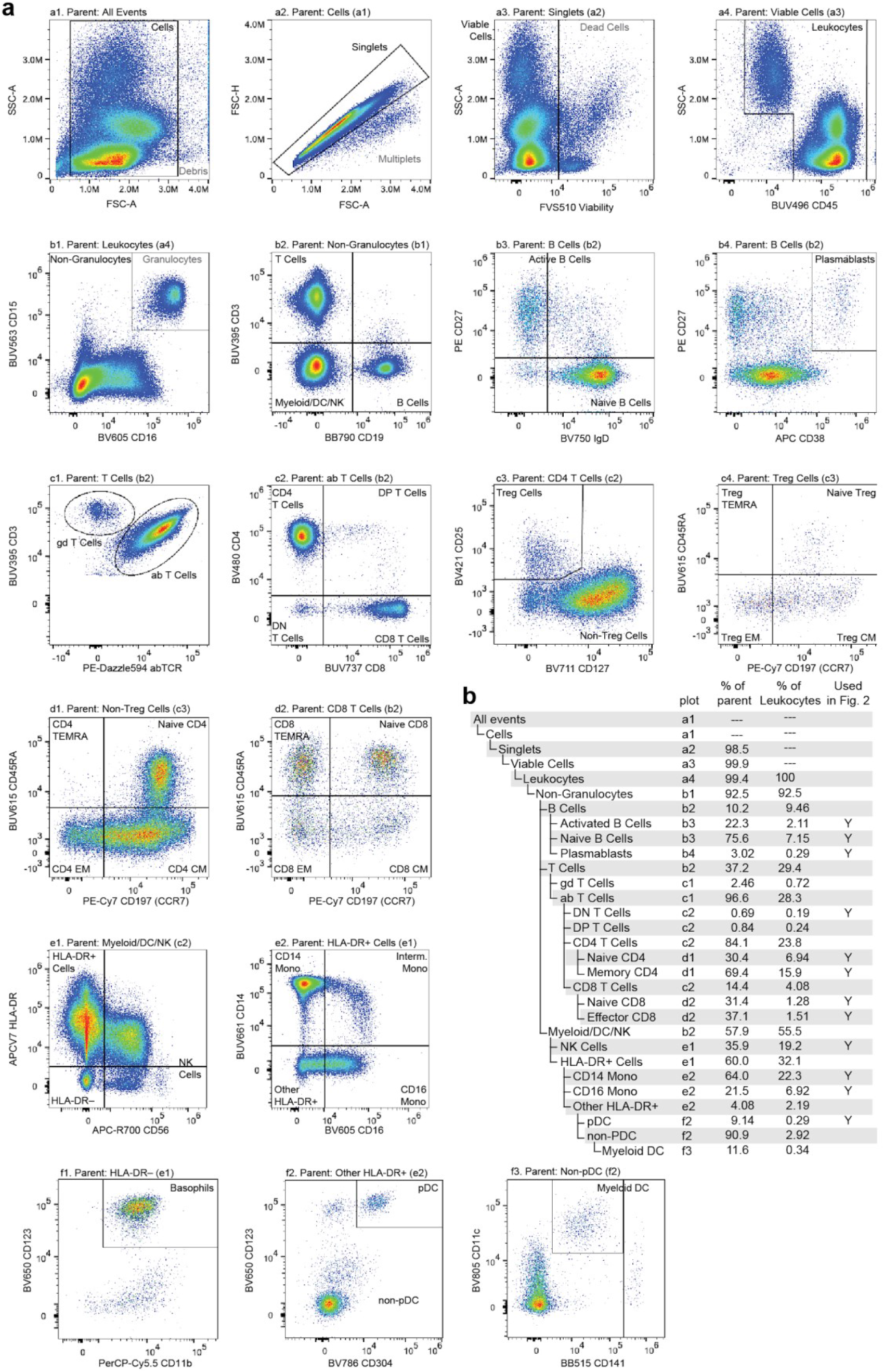
**a**, Flow cytometry gating used to classify and quantitate cell type abundance. Cells were stained with a panel of 25 fluorescently labeled antibodies in total, fluorescence was measured using a Cytek Aurora spectral flow cytometer, and events were manually gated to assess cell type abundance. Pairwise feature plots show the gates used for cell type assessment. Plots are arranged in rows (labeled a-f) and columns (labeled 1-4). The plot and label of the parent gate is at the top of each plot, and gates derived from each set of markers are labeled within the plots. **b**, Gating hierarchy used to quantify cell types, with references to the plot in which each gate is defined. Cell type labels used to generate reference proportions in **Figure 2** are specified in the last column.

**Figure 3 – figure supplement 1.**
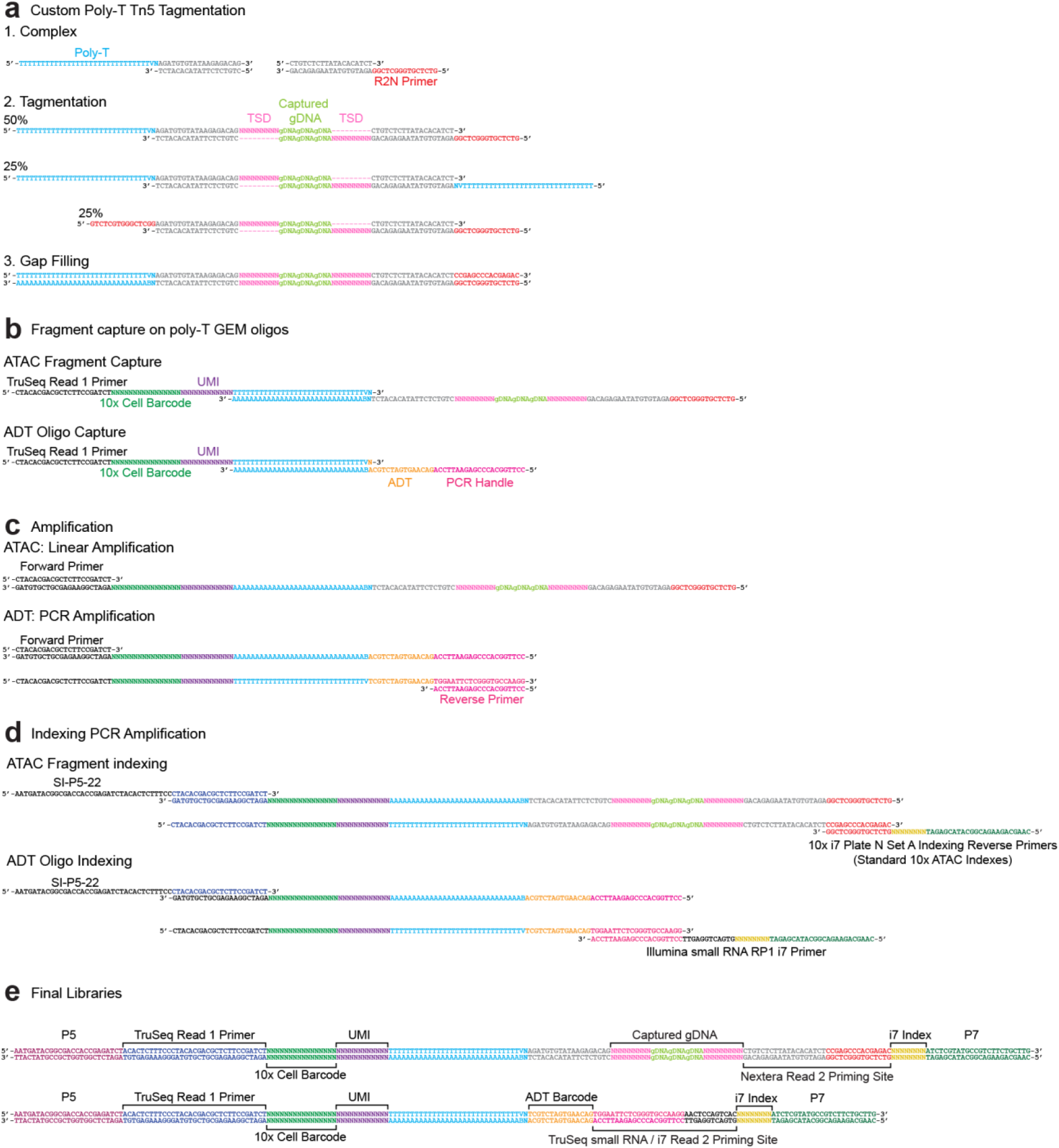
Sequence-level ICICLE-seq design and workflow. **a**, Tn5 transposase complexes are formed by annealing Poly-T and Nextera R2N top oligos to MOSAIC-end bottom oligos to form double stranded DNA complexes. Complexes are bound to Tn5 transposase in the mosaic-end regions and transposed into target gDNA via a tagmentation reaction. Approximately half of the resulting ATAC fragments have both a Poly-T and R2N overhang and are capable of being barcoded and indexed in subsequent amplification reactions. **b**, Following GEM generation and gap-filling, DNA fragments are denatured to allow annealing to 10x Genomics 3’ bead oligos and extension by a DNA polymerase, resulting in the addition of cell barcode and UMI sequences. **c**, Barcoded ATAC fragments are amplified linearly using a forward primer, while barcoded ADT fragments are amplified exponentially by PCR through the addition of an ADT specific reverse primer. **d**, Intermediate libraries are amplified and Illumina P5 and indexed P7 adapter sequences are added by PCR. Prior to amplification, libraries are divided into separate ATAC and ADT reactions and amplified using library specific P7 primers. **e**, Final ATAC and ADT library structure includes all components required for sequencing. The TruSeq Read 1 Primer reads the 10x Cell Barcode and UMI for both libraries as Read 1, followed by readout of the i7 Index sequence with the Nextera Read 2 Priming Site (ATAC) or TruSeq small RNA /i7 Read 2 Priming site (ADT). After strand switching, these two latter binding sites are used to read out the captured gDNA sequence (ATAC) or ADT Barcode as Read 2.

**Figure 3 – figure supplement 2.**
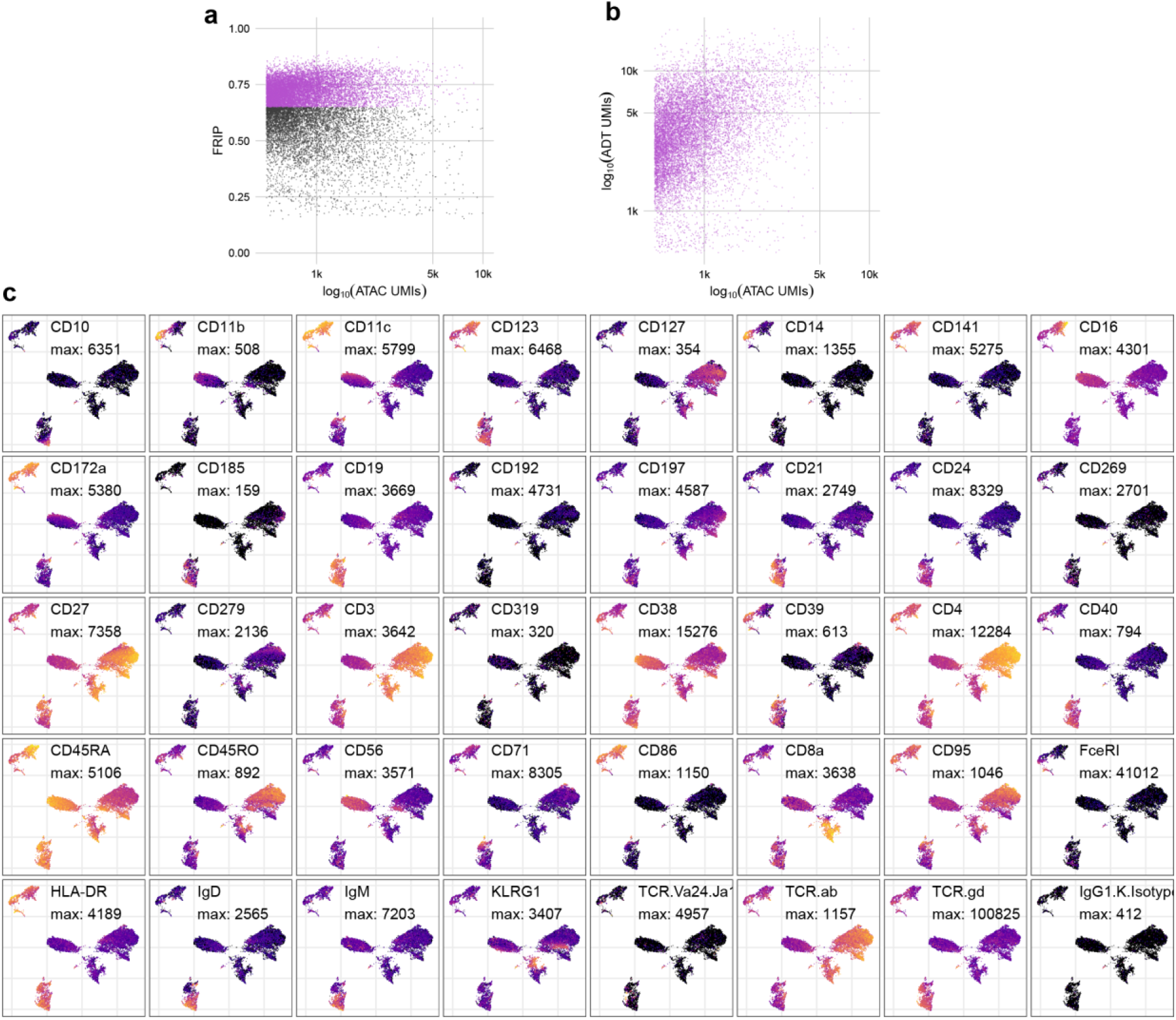
**a**, QC metrics plot for the ATAC-seq component of ICICLE-seq. Cell barcodes with FRIP > 0.65 (purple points) were selected for downstream analysis. **b**, Comparison of the number of ATAC UMIs (x-axis) to ADT UMIs (y-axis) for each cell barcode that passed the QC cutoff in (**a**). **c**, UMAP plots based on ADT UMI counts. Each panel shows log-transformed expression of a different marker, and values scaled between 0 (black) and the maximum observed value (yellow).

**Figure 4 – figure supplement 1.**
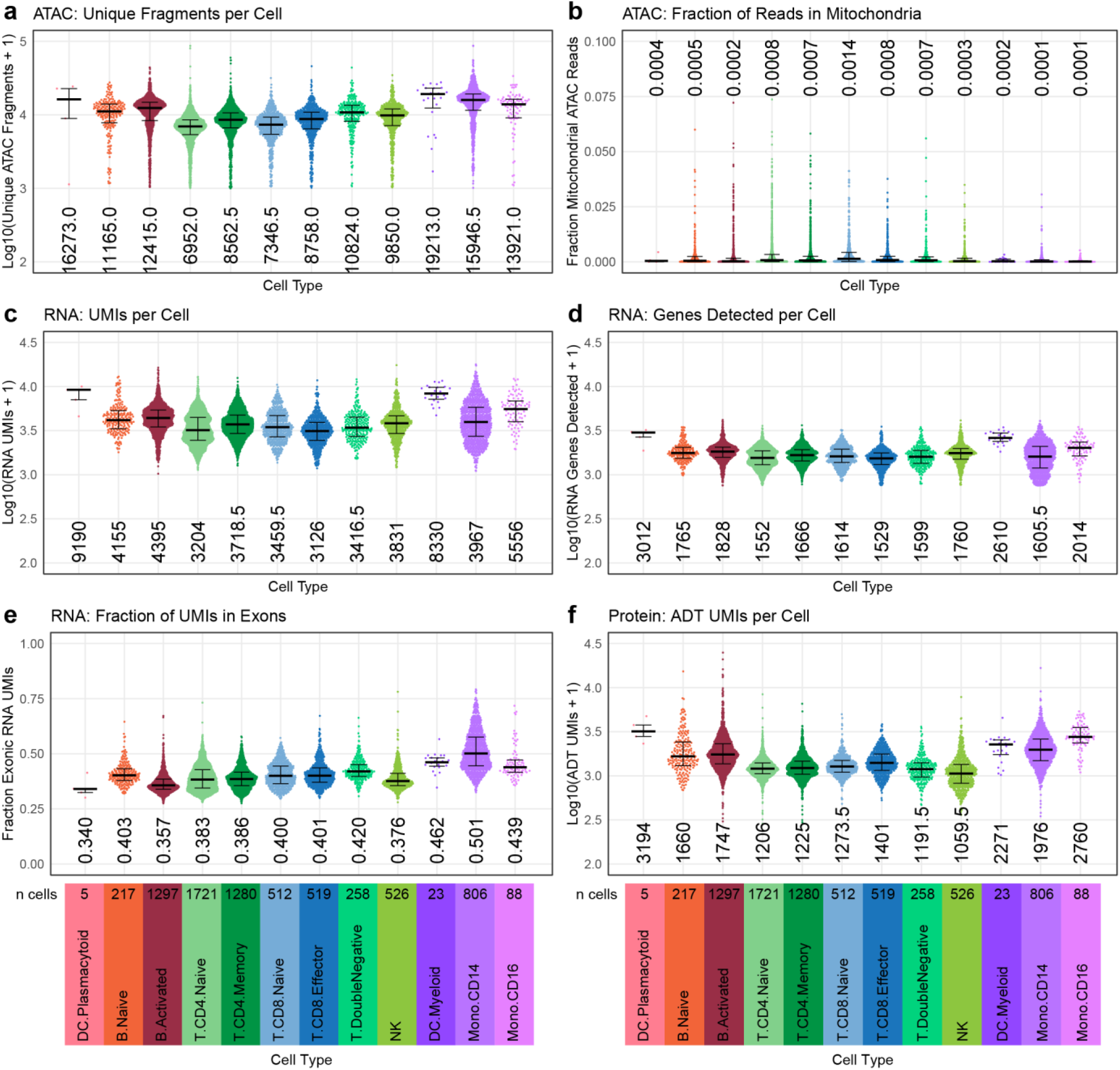
Quasirandom-jittered plots (jittered only on x-axis) showing various QC metrics from TEA-seq cells that passed all QC criteria (**Methods**). In each panel, cells are separated by cell type (x-axis). Median values per cell type for each metric are printed within the plot region at the x-axis position corresponding to each cell type. Types and number of cells in each category are displayed below the bottom row and apply to all plots in each column. **a**, Number of unique ATAC fragments detected per cell (y-axis, Log10 scale). **b**, Fraction of raw ATAC fragments that were aligned to mitochondrial regions (y-axis, linear scale, max = 0.1). **c**, Number of RNA UMIs assigned to each cell (y-axis, Log10 scale). **d**, Number of genes detected by RNA-seq for each cell (y-axis, Log10 scale). **e**, Fraction of RNA UMIs from exonic regions (y-axis, linear scale). **f**, Number of ADT UMIs assigned to each cell (y-axis, Log10 scale).

**Figure 4 – figure supplement 2.**
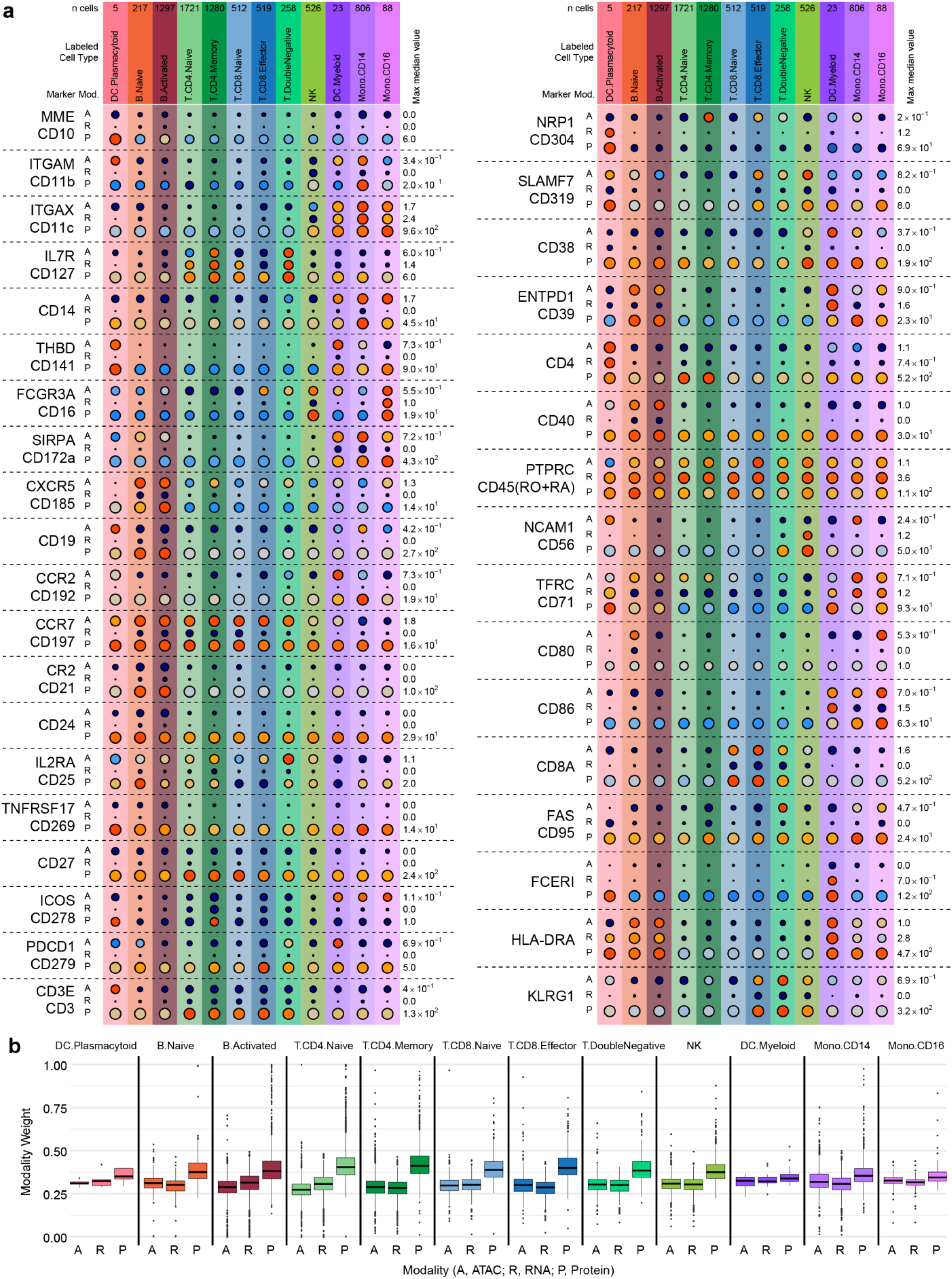
**a**, Detection of 36 protein markers with corresponding scATAC-seq GeneScores and RNA UMIs from TEA-seq cells. Markers included in the ADT set (**Supplemental Table 6**) were filtered to retain only those with both corresponding GeneScores and RNA UMI counts. Each horizontal section of the plot, separated by dashed lines, presents detection for a single marker. When gene symbols differ from the protein marker name, the gene name is shown above the antibody name. Each section is subdivided into rows for each of the 3 assays (A, ATAC; R, RNA; P, Protein). The size of each point represents the fraction of cells within each cell type (columns) with > 0 detection for each marker within each modality (larger points = greater fraction). The color of each point represents the median of the detected value for each assay, normalized within each row between zero (dark blue) and the maximum value for each feature and modality (provided at the right of each row). For the comparison to PTPRC/CD45, the ADT counts for CD45RA and CD45RO were summed. Mod, modality. **b**, Weight contributions of each modality to the WNN graph used to generate the UMAP in panel g. Boxplots represent the modality weight distribution of individual cells within each cell type; Heavy lines mark the median value, box boundaries represent the 25th and 75th quantiles, and whiskers extend to 1.5 times the interquartile range above the box boundaries.

### Supplementary Tables

**Supplementary Table 1.** Antibody-fluorophore conjugates used to characterize PBMC cell type abundance by flow cytometry before and after anti-CD15 bead-based neutrophil depletion.

**Supplementary Table 2.** Quantification of major cell populations by flow cytometry before (Pre-depletion) and after (Post-depletion) anti-CD15 bead-based neutrophil depletion of either Ficoll-purified or Leukapheresis-purified PBMCs. The gating strategy used to assess these populations is presented in **Figure 1 – figure supplement 4**.

**Supplementary Table 3.** Antibody-fluorophore conjugates used to assess PBMC cell type populations for evaluation of cell type labeling.

**Supplementary Table 4.** Quantification of PBMC cell type populations using the antibody panel provided in **Supplementary Table 3**. The gating strategy used to assess these populations is presented in **Figure 2 – figure supplement 1**. Type labels and proportions used for comparisons of scATAC-seq cell type labeling in **Figure 2d** are provided in the last two columns. When two gated populations are combined to tabulate a single cell type, the same cell type label is listed beside each gate. Cell type proportions are listed only at the first instance of each cell type.

**Supplementary Table 5**. Custom oligonucleotide sequences used to generate ICICLE-seq libraries.

**Supplementary Table 6.** Antibody-oligo conjugates (BioLegend TotalSeq-A) used for ICICLE-seq and TEA-seq experiments to label PBMC cell types.

**Supplementary Table 7.** Sources and presentation of human biological specimens utilized in this study.

